# Shared latent representations of speech production for cross-patient speech decoding

**DOI:** 10.1101/2025.08.21.671516

**Authors:** Z. Spalding, S. Duraivel, S. Rahimpour, C. Wang, K. Barth, C. Schmitz, S. P. Lad, A. H. Friedman, D. G. Southwell, J. Viventi, G. B. Cogan

## Abstract

Speech brain-computer interfaces (BCIs) can restore communication in individuals with neuromotor disorders who are unable to speak. However, current speech BCIs limit patient usability and successful deployment by requiring large volumes of patient-specific data collected over long periods of time. A promising solution to facilitate usability and accelerate their successful deployment is to combine data from multiple patients. This has proven difficult, however, due to differences in user neuroanatomy, varied placement of electrode arrays, and sparse sampling of targeted anatomy. Here, by aligning patient-specific neural data to a shared latent space, we show that speech BCIs can be trained on data combined across patients. Using canonical correlation analysis and high-density micro-electrocorticography (μECoG), we uncovered shared neural latent dynamics with preserved micro-scale speech information. This approach enabled cross-patient decoding models to achieve improved performance relative to patient-specific models facilitated by the high resolution and broad coverage of μECoG. Our findings support future speech BCIs that are more accurate and rapidly deployable, ultimately improving the quality of life for people with impaired communication from neuromotor disorders.

## Main

Speech is an intricate coordination of muscle movements that shape sounds into words^1^. Paralysis of these muscles by neuromotor disorders such as amyotrophic lateral sclerosis (ALS) and brainstem stroke can disrupt speech, resulting in debilitating losses to communicative ability and a reduced quality of life^2–6^. Speech brain-computer interfaces (BCIs) have shown promise in restoring communication to individuals with neuromotor disorders by using machine learning models to accurately decode intended speech units directly from brain activity^7–10^.

Speech BCIs can be broadly divided into non-invasive approaches, such as electroencephalography (EEG), and invasive approaches, such as electrocorticography (ECoG). Non-invasive BCIs offer advantages in safety and ease of deployment but are limited by lower spatial resolution and susceptibility to noise, which can constrain their ability to resolve fine-grained articulatory representations required for high-accuracy speech decoding^11,12^. In contrast, invasive BCIs provide higher spatial and temporal resolution recordings directly from cortical populations, enabling more accurate decoding of speech-related neural activity^7–10^, but introduce challenges related to surgical implantation, limited and variable electrode coverage, and patient-specific sampling of neural populations. Invasive recording modalities have become a primary focus for achieving high-accuracy speech decoding despite these challenges.

A key limitation of invasive speech BCIs is their reliance on patient-specific approaches that require resource-intensive, individualized data acquisition from each patient – typically spanning multiple weeks – before their associated machine learning models can be trained to achieve maximal accuracy^8^. This approach is taken to mitigate the effects of varied sensor placement, sparse sampling of the targeted anatomy, and neuroanatomical variability across patients^13^. These long training and deployment times make patient use cumbersome and hinder their adoption as assistive communication devices. Future speech BCIs that successfully combine data across multiple patients may accelerate training by requiring minimal data collection from new patients. Such cross-patient speech BCIs could be rapidly deployable and demonstrate greater robustness across a broader patient population.

For successful cross-patient speech decoding, there must be similarity in the neural representations of speech production across patient recordings. Invasive speech decoding studies typically record from the sensorimotor cortex (SMC)^14–16^ with intracranial electroencephalographic devices, such as ECoG^17–20^ or intracortical microelectrode arrays^21,22^, from which sampled representations of speech production may differ from patient to patient due to variability in electrode array placement or individual neuroanatomy. However, recent work has demonstrated that for motor activity, there is a low-dimensional structure in the time-varying neural population activity of the motor cortex^23–25^, known as latent dynamics, that is shared across individuals. For example, in non-human primates, the latent dynamics of individual subjects performing limb movement can be aligned and pooled across animals to train a cross-animal movement decoding model^26^. We hypothesize that this phenomenon extends to human speech production. That is, the latent dynamics underlying the motor control of articulator muscles in the vocal tract to produce speech have a shared representation across separate patients despite variability in electrode placement and individual neuroanatomy (Fig. 1a). These latent dynamics can be aligned and pooled across patients to enable training of cross-patient speech BCIs.

**Figure 1.**
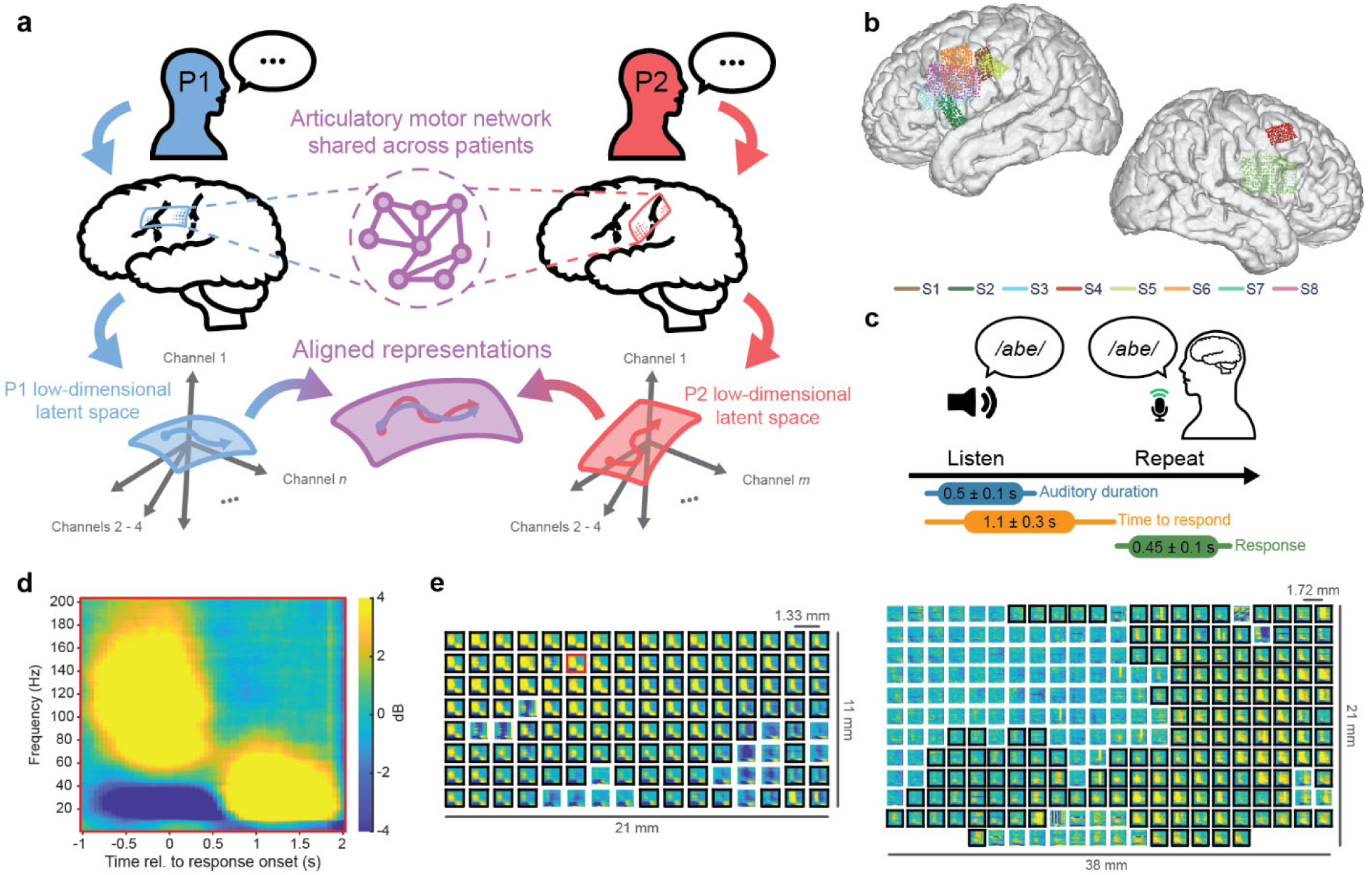
Shared latent dynamics underlying speech production. **a.** Despite differences in electrode placement and individual neuroanatomy, we propose that the neural population activity that controls articulatory muscles for speech production is shared across separate patients and can be elucidated by aligning their latent dynamics to a common space. **b.** Neuroanatomical location of μECoG arrays for all patients projected onto a common brain (MNI-152, see Fig. S1 for patient-specific array locations). **c.** Speech repetition task. Patients are presented with an auditory non-word stimulus that is composed of three phonemes (e.g. */abe/*) and are instructed to immediately repeat this phoneme sequence. Timing shows statistics of task performance. For information on experimental design see Methods – *Speech task* **d.** Example spectrogram showing high-gamma (HG; 70-150 Hz) power centered around the response onset (*t* = 0). **e.** Spectrograms across 128-channel (left, S1) and 256-channel (right, S3) μECoG arrays. Black borders indicate channels with significant HG power during patient response relative to a pre-stimulus baseline (FDR-corrected non-parametric permutation tests). The red border indicates the example spectrogram from **d**. To identify electrodes with significant HG activation during patient responses, we compared the average (over time) HG power around the patient response onset (-250 ms to 250 ms) to a pre-stimulus baseline period (-500 ms to 0 ms relative to stimulus presentation) for each electrode using a FDR-corrected 10,000-iteration one-sided permutation test.

We additionally hypothesize that the elucidation of shared latent dynamics requires an accurate sampling of speech-relevant neural population activity. We have previously shown that accessing the rich spatiotemporal representations of speech production in the SMC requires precise neural interfaces with high spatial-specificity and broad coverage. This dense, broad sampling of the cortex has been achieved using micro-electrocorticographic (μECoG) arrays to record from a large surface of the brain with a high density of micro-contacts^27^, which has allowed resolution of micro-scale features for both epilepsy^28^ and speech production^29^. While surface recording via μECoG relies on activity in neural local field potentials (LFPs) and cannot access the same multi-unit activity that previous studies used to investigate these cross-patient motor latent dynamics^26^, previous work has shown strong similarity between latent dynamics extracted from high-frequency LFP bands and multi-unit sources^30^. Together, these results suggest that μECoG has sufficient spatial-specificity, coverage, and signal quality to enable successful cross-patient alignment for speech BCIs.

Here, we investigate cross-patient representations of speech production by performing acute neural recordings with high-density μECoG arrays over the SMC of eight patients during a speech repetition task. We use dimensionality reduction techniques to reduce high channel-count μECoG recordings into low-dimensional latent dynamics, and we subsequently learn linear transformations that align these latent dynamics across patients. We show that cross-patient latent dynamics encode speech information relevant to the control of articulator muscles in the vocal tract. These shared speech encodings enable the training of cross-patient speech decoding models, which in turn outperform patient-specific decoding models. Lastly, we show that micro-scale sampling of the SMC with broad coverage is critical for learning the proper alignment between patients that preserves speech information.

Our results demonstrate improved speech decoding for BCIs facilitated by a shared, cross-patient representation of speech production encoded in the latent dynamics representing SMC population activity. These findings enable future speech BCIs with accelerated deployment times and increased generalizability, resulting in improved quality of life for individuals who have lost the ability to speak due to neuromotor disorders.

## Results

We recorded speech-relevant neural activity from the surface of the SMC using high-density, high channel-count μECoG arrays in eight patients undergoing awake neurosurgical procedures (four with 128 channels – 1.33 mm spacing, four with 256 channels – 1.72 mm spacing, see Table S1 for summary of patient information). Fig. 1b shows the electrode array locations of all patients projected to an average brain (see Fig. S1 for array locations on patient-specific brains). Recordings were performed intraoperatively while patients were instructed to repeat non-word sequences of three phonemes, or sound units (e.g. */b/* in bat), immediately after hearing an auditory presentation of the non-word (Fig. 1c, see Methods and Table S2). Using spectrotemporal analysis of the recorded neural signals, we observed neural activations around patient response onset (Fig. 1d). Specifically, we found strong increases in power in the high-gamma (HG; 70-150 Hz) band, which has been shown to be a robust marker of speech production in the SMC^8,19,31^ that correlates well with multi-unit neural firing^32–34^. Significant HG activations during patient responses (relative to a pre-stimulus baseline, FDR-corrected non-parametric permutation test – see Methods) were observed in many electrodes across the entirety of each μECoG array (significant electrode amounts in Table S3), exhibiting unique spatial activation patterns (Fig. 1e, all patients in Fig. S3).

### Alignment of cross-patient latent dynamics

Speech trials for each patient were grouped into four categories representing types of articulation corresponding to the phonemes presented in our stimuli (low: */a/, /e/*, high: */i/, /u/*, labial: */p/, /b/, /v/,* dorsal: */k/, /g/*). For simplicity, we classified trials by the articulator type of the first phoneme in the spoken sequence, resulting in 12 “low” non-words, 14 “high” non-words, 14 “labial” non-words, and 12 “dorsal” non-words. There were distinct spatiotemporal patterns of HG activity specific to each type of articulation (Fig. 2a, all patients in Fig. S4). While patterns differ across articulators, they also differ across patients. This can be seen across all patients, but we illustrate this difference for two representative patients (S1 and S2 – Fig. 2a). These patient-specific differences in spatiotemporal speech representations demonstrate the necessity of specialized alignment techniques in uncovering a shared representation of speech across patients.

**Figure 2.**
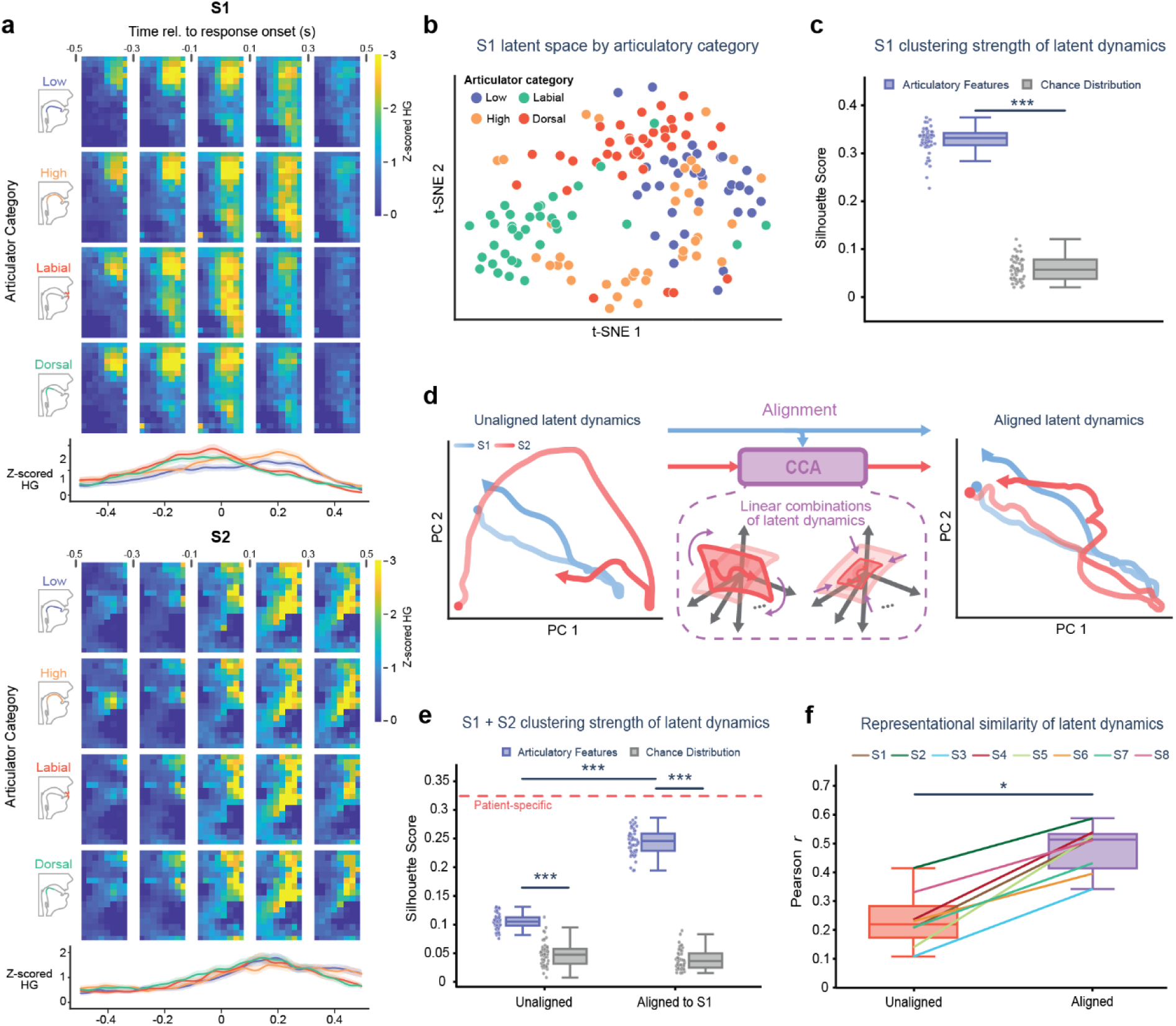
Alignment of latent dynamics across patients. **a.** Spatiotemporal activation patterns of HG neural activity (z-scored relative to pre-stimulus baseline) across the μECoG arrays for S1 and S2 grouped by different articulatory gestures. Vocal tract diagrams show visual representations of articulatory category (low, high, labial, dorsal). HG traces below the heatmaps show average activity across all channels for each articulator category. The spatiotemporal HG patterns of these articulatory gestures are distinct across patients, motivating a need for alignment for cross-patient grouping. **b.** Latent dynamics are extracted from HG activity using principal component analysis (PCA). Projection of S1 latent dynamics to a 2D space as shown by t-distributed stochastic neighbor embedding (t-SNE) which qualitatively demonstrates the clustering of latent dynamics by articulator type. **c.** Quantification of articulatory clustering strength in **b** by the silhouette score of points in t-SNE space. The silhouette score for latent dynamics is significantly higher than that of a chance distribution generated by shuffling the ground truth articulator labels (\*\*\**p < 0.001*, Mann-Whitney U test). **d.** Latent dynamical trajectories for S1 and S2, as visualized in 2D PC space. The trajectories appear dissimilar. Alignment of S2 latent dynamics to S1 is performed with canonical correlation analysis (CCA). Resultant aligned latent dynamics appear more visually similar. **e.** As in **c**, but with data from both S1 and S2. Without alignment, the latent dynamics articulatory clustering strength is above chance (*p < 0.001*, Mann-Whitney U test) but much lower than the patient-specific S1 mean. With alignment, the articulatory clustering strength is significantly higher than both the unaligned case (*p < 0.001*, Mann-Whitney U test) and chance (*p < 0.001*, Mann-Whitney U test), and much closer to the patient-specific S1 mean. **f,** Representational similarity analysis on cross-patient latent dynamics. Across all patients, cross-patient similarity in speech information (Pearson *r* between representational dissimilarity matrices) represented by latent dynamics are significantly higher when latent dynamics are first aligned with CCA alignment instead of left unaligned (\**p < 0.05*, Wilcoxon signed-rank test). For latent dynamics clustering strength (quantified by the silhouette score), comparisons between distributions (articulatory features vs chance) were performed with Mann-Whitney U (MWU) test. MWU tests were also used for cluster strength comparisons when combining data from both S1 and S2, where FDR correction was applied. Differences in paired distributions for cross-patient representational similarity of latent dynamics were compared with a Wilcoxon signed-rank test.

To extract latent dynamics from these high-dimensional data, we use principal component analysis (PCA) as a linear dimensionality reduction tool to decompose HG activity (± 500 ms around speech onset) across the significant channels into a low-dimensional set of principal components (PCs) while preserving timing information. Articulatory information encoded in the latent dynamics can be visualized by further projecting the latent dynamics into a two-dimensional space using t-distributed Stochastic Neighbor Embedding (t-SNE, see Methods), where separation between clusters for different articulator types was qualitatively apparent (shown here for S1 – Fig. 2b). Clustering strength of the latent dynamics in this space was quantified using the silhouette score (for additional clustering strength metrics, see Fig. S5), which showed significantly better separation of articulatory feature clusters than in a chance distribution where ground-truth cluster identities were shuffled (Fig. 2c; Mann-Whitney U test, *U =* 2500.0*, n_1_ = n_2_ = 50, p* =7.07e-18). This shows that PCA successfully extracts latent dynamics that are informative to speech.

The latent dynamics showed distinct trajectories through their respective latent spaces, as visualized in two-dimensional PC space (S1 and S2: Fig. 2d, left). We sought to functionally align these latent dynamics to a similar space to train speech decoding models that learn a similar representation of speech production from multiple patient sources. To model this functional alignment, we used canonical correlation analysis (CCA) to identify linear combinations of patient-specific latent dynamics such that they are maximally correlated across patients. This resulted in patient-aligned latent dynamics that show more similar trajectories through a common latent space (shown for S1 and S2: Fig. 2d, right). Repeating our previous technique of clustering strength on latent dynamics, we found weaker across-patient clustering when these latent dynamics were unaligned relative to patient-specific clustering, indicating differences in articulator representations across patients (Fig. 2e). With alignment, clustering strength was significantly higher than when unaligned (Mann-Whitney U test, *U =* 2500*, n_1_ = n_2_ =* 50, FDR-corrected *p* = 1.06e-17) and much closer to patient-specific values. As t-SNE operates unsupervised, it is decoupled from any potential leakage of label information that could occur during alignment. Therefore, silhouette score quantification of clustering in the t-SNE space of aligned latent dynamics represents a valid quantification of aligned cross-patient clustering, and shows that CCA alignment can successfully map latent dynamical representations of speech across different patients.

To further examine the ability of CCA to align representations of speech across all patients, we used representational similarity analysis (RSA) to quantify cross-patient similarity in latent dynamical representations of articulatory information. Specifically, we measured the Pearson correlation between representational dissimilarity matrices (RDMs) constructed from cross-patient latent dynamics with and without alignment (Fig. S6; see Methods). To better capture speech information and task structure in these RDMs, we grouped trials by their full, three-length articulator sequence (e.g. “labial-low-dorsal”) instead of simply by the first articulator in the spoken sequence. Across all patients, we found that correlation between RDMs significantly improved when latent dynamics were aligned to a common patient space prior to RDM construction instead of unaligned in separate patient spaces (Fig. 2f; Wilcoxon signed-rank test, *W* = 0, *n =* 8, *p* = 0.01). Additionally, we found that alignment does not trivially result in an increased correlation between articulatory RDMs, as RDMs constructed from aligned latent dynamics with permuted articulatory labels (and thus no relevant organization of articulatory information) showed significantly lower correlations than those constructed with ground truth articulatory labels (Fig. S7; Wilcoxon signed-rank test, *W* = 0, *n =* 8, FDR-corrected *p* = 0.01). To further show that speech-relevant neural activity is important to cross-patient alignment, we also quantified articulatory representational similarity in a control condition using surrogate data. Specifically, we replaced the true neural data with surrogate data generated by the Tensor Maximum Entropy (TME) method, which is randomly generated with the constraint of preserving the first and second order statistical structure of our neural data (equal mean and covariance across trials, channels, and time; see Methods). As such, this surrogate data is random with respect to the speech sequences spoken during trials and should not encode any relevant articulatory information. Cross-patient representational similarity from this surrogate data control was significantly lower than that of our original aligned latent dynamics (Fig. S7; Wilcoxon signed-rank test, *W* = 0, *n =* 8, FDR-corrected *p* = 0.01), indicating that speech information encoded in the true neural activity is important to successful cross-patient alignment. These results demonstrate the success of CCA in aligning latent dynamics across patients to elucidate a shared representation of speech production.

### Cross-patient alignment preserves spatial articulator maps

Previous work leveraging the high spatial resolution of μECoG showed micro-scale patterns of spatial tuning to different articulator types^29^. As the alignment of latent dynamics showed that articulator representations could be co-aligned temporally across patients, we hypothesized that spatial tuning to articulatory features would also be preserved following alignment. To investigate this, we first calculated single-electrode tuning to the four articulator categories. For each electrode, we trained a linear discriminant analysis decoding model to classify articulator type. We quantified the tuning of an electrode to an articulator by its receiver-operator-characteristic area-under-the-curve (ROC-AUC) value. Fig. 3a shows the spatial arrangement of electrode-articulator tunings for S1, where distinct spatial maps for each articulator type are visible.

**Figure 3.**
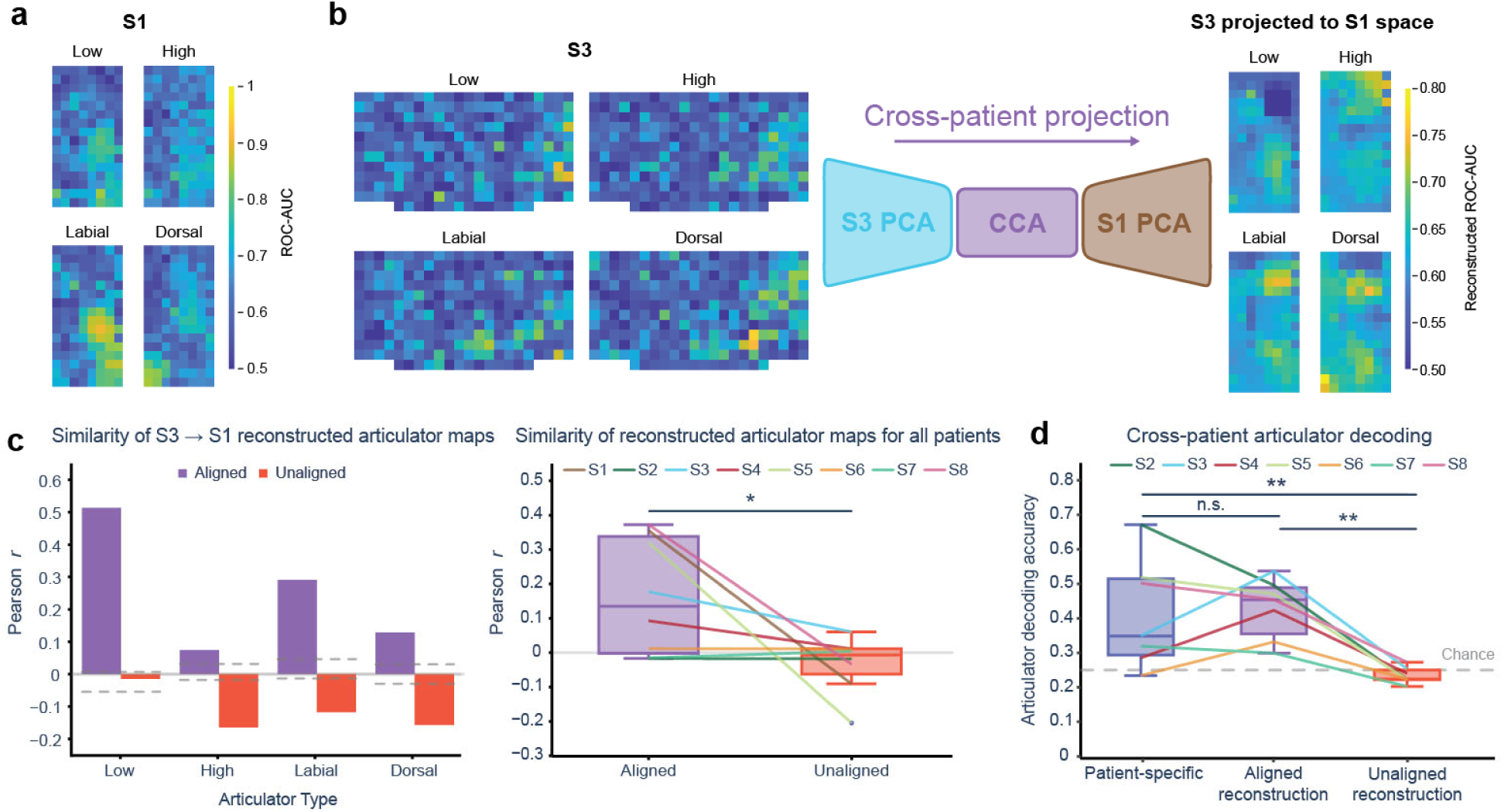
Cross-patient alignment preserves spatial representations of articulatory features. **a.** Spatial map of electrode tuning to different articulator categories for S1 (128 channels), as quantified by the ROC-AUC score from univariate electrode decoding. Different articulators have distinct spatial representations over the array. **b.** Same as in **a**, but for patient S3 (256 channels). Cross-patient projection enabled reconstruction of a source patient’s (S3) articulator tuning in a target patient’s (S1) space. **c.** (Left) Quantification of Pearson correlation between reconstructed articulator maps from source patient S3 and ground-truth articulator maps for target patient S1. Reconstructed maps are more similar when using CCA alignment than without alignment across all articulator types. Horizontal dashed lines indicate an interval of trivial correlation values, represented by 95% confidence intervals on correlation values from a null distribution of reconstructions performed with permuted CCA weights (*n* = 100). Correlation values above this interval suggest non-trivial alignment to the target articulator map, while correlation values below this interval suggest misalignment, or non-trivial anti-correlation. (Right) Similarity in articulator map reconstructions for all target patients (*n* = 8, each dot is a target patient). For each target patient, reconstructed maps were averaged across all source patients within articulator type to produce a resultant map to compare against the ground-truth articulator map. Reported similarity values for each target patient are the average similarity across all four articulator types. Similarity in reconstructed maps from aligned latent dynamics are significantly higher than those without alignment (\*\*\**p < 0.001*, Wilcoxon signed-rank test). **d.** Articulator decoding from cross-patient data. Support vector machine (SVM) decoding models are trained to classify articulator type from latent dynamics. There is no significant difference in accuracy between patient-specific SVMs and SVMs trained on S1’s data then tested on data from other patients that have been aligned and reconstructed to S1’s space (*p = 0.79*, exact two-sided paired permutation test). Without alignment, cross-patient decoding accuracy is significantly lower (\*\**p < 0.01*, exact two-sided paired permutation test). Reconstructed articulator map similarity was compared between aligned and unaligned contexts with a Wilcoxon signed-rank test. Cross-patient articulator decoding accuracy distributions were compared with exact two-sided, paired permutation tests and FDR correction. Comparison of cross-patient articulator decoding accuracies to chance accuracy was performed using Wilcoxon signed-rank tests corrected with FDR correction.

To investigate the cross-patient preservation of these maps, we used CCA alignment to project spatial articulator information from one patient’s space to another. We first used PCA to extract latent dynamics from two patients and learned a CCA transformation to align them. The transformations learned in this process enable the projection of data from a source patient’s “electrode-space” to a target patient’s electrode-space, which we call cross-patient projection. Specifically, electrode-level data can be projected to the source patient’s latent space with the source patient’s PCA transform, (optionally) aligned to the target patient’s latent space, and then projected to the target patient’s electrode-space by applying the inverse of target patient’s PCA transform. We illustrate this analysis with S3 as the source patient and S1 as the target patient (Fig. 3b). After learning the PCA and CCA transformations, the electrode-articulator tunings for the source patient (Fig. 3b, left) were projected to the target patient’s space. We denote these projected spatial maps as reconstructed articulator maps (Fig. 3b, right). The Pearson correlation between ground-truth articulator maps and reconstructed articulator maps was calculated to quantify similarity in the reconstructed spatial tunings. We found that the similarity between reconstructed and target articulator maps was higher when cross-patient projection was performed with alignment than without (Fig. 3c, left). Additionally, we found that the target patient’s articulator maps could be reconstructed with high similarity for certain articulator categories (low, labial, dorsal) by averaging reconstructed maps across all source patients (Fig. S8 – illustrated for target patient S1), showing that articulator information in one patient is well-represented across multiple patient sources. Across all target patients, we found significantly higher similarity between reconstructed and ground-truth maps with alignment (Fig. 3c, right) than without (Wilcoxon signed-rank test, *W =* 3, *n =* 8, *p* = 0.04), indicating that cross-patient alignment results in improved preservation of spatial articulator tuning. To confirm this similarity between ground-truth and reconstructions of articulator maps with alignment was indeed driven by shared speech information encoded in the recorded neural activity, we performed a control analysis again using TME-generated surrogate data in place of our original neural data (Fig. S9). When decomposition and alignment transformations were learned using this surrogate data that is random with respect to speech information, we found reduced mean correlations between ground truth and reconstructed articulator maps with alignment across all patients and articulators (Fig. S9a; mean: surrogate aligned = -0.09, aligned = 0.16), though with no significant effect (FDR-corrected *p =* 0.07, exact two-sided paired permutation test), which may be due to our limited number of patients.

We validated the preservation of articulatory information across patients by training support vector machine (SVM) decoding models to classify articulatory features from patients that had not been used for training. Specifically, we trained a singular SVM on HG signals (± 500 ms around speech onset) from one patient and evaluated its accuracy when tested on all other patients projected to the single patient’s space, both with alignment and without (Fig. 3d – illustrated for S1). As a baseline, we compared cross-patient accuracies to the accuracy of patient-specific SVMs for each patient. We found both patient-specific and cross-patient decoding on aligned reconstructed data yielded above chance articulator decoding accuracies (chance = 0.25; mean: patient-specific = 0.39, Wilcoxon signed-rank test, W = 1, *n =* 7, *p* = 0.047; mean: aligned reconstruction = 0.43, Wilcoxon signed-rank test, W = 0, *n =* 7, FDR-corrected *p* = 0.047). Additionally, we found no significant difference between patient-specific and aligned cross-patient decoding accuracies (*n =* 7, FDR-corrected *p =* 0.79, exact two-sided paired permutation test), indicating that the aligned cross-patient SVM performs just as well as the patient-specific SVM despite the separate patient sources. Without alignment, there was a significant decrease in cross-patient decoding accuracy relative to the aligned context (mean unaligned reconstruction = 0.23; *n =* 7, FDR-corrected *p =* 0.002, exact two-sided paired permutation test), which was expected due to the dissimilarity found between reconstructed articulator maps without alignment and ground-truth articulator maps. There was no significant difference between unaligned cross-patient decoding accuracies and chance accuracy (chance = 0.25; Wilcoxon signed-rank test, W = 4, *n =* 7, FDR-corrected *p* = 0.11), demonstrating the necessity of alignment in elucidating shared neural representations of speech. We repeated this aligned decoding analysis with our TME-generated surrogate data to again confirm that success of the aligned reconstructions was driven by speech information encoded in the original neural data (Fig. S9b). We observed a significant decrease in cross-patient decoding accuracies on aligned reconstructed data in the surrogate control relative to that of our original data (mean: surrogate aligned reconstruction = 0.27, aligned reconstruction = 0.43; *n =* 7, FDR-corrected *p =* 0.01, exact two-sided paired permutation test). Overall, we have demonstrated that cross-patient alignment preserves important spatial characteristics of articulatory features in the SMC, enabling successful cross-patient speech decoding.

### Expanded cross-patient datasets improve speech decoding accuracy

Given the preservation of speech information in aligned, cross-patient latent dynamics, we investigated if cross-patient datasets could be used to address data limitations in speech BCI training and improve their performance. With a selected target patient, additional data from other patients can be aligned to this target patient and appended to the target datasets, enabling vastly expanded training datasets relative to patient-specific contexts. To test this, we trained SVM models on aligned cross-patient latent dynamics during speech (total combined dataset size across all patients: ∼8.2 minutes utterance duration, ∼68.5 minutes experiment duration, see Table S3 for amount of speech data per patient) and we evaluated their accuracy on a held-out set from the target patient using 20-fold cross-validation. We then compared these results to patient-specific latent dynamics and unaligned cross-patient latent dynamics. While thus far, our analyses focused on decoding articulatory features, here we evaluated the accuracy of classifying distinct phoneme units present within these articulator categories (e.g. /k/, /a/, /b/) and in all three position of the spoken phoneme sequence to mimic translational BCI use cases where phonemes are rapidly decoded and strung together into words and sentences.

We found significant differences in decoding accuracy across patient-specific, unaligned cross-patient, and aligned cross-patient decoding contexts (Fig. 4a; 9-way phoneme decoder, chance = 0.11; repeated measures ANOVA, *F_2,14_* = 12.59, *p =* 0.007). In decoding models trained on unaligned cross-patient latent dynamics, individual phoneme decoding accuracy was significantly lower relative to patient-specific decoding (unaligned cross-patient: mean = 0.19, 95% CI = [0.15, 0.24]; patient-specific: mean = 0.24, 95% CI = [0.17, 0.31]; follow-up FDR-corrected paired t-test, corrected *p =* 0.03; effect size *d* = 0.57). Models trained on aligned cross-patient latent dynamics, however, exhibited strong phoneme decoding accuracy – even outperforming patient-specific models (aligned cross-patient: mean = 0.31, 95% CI = [0.22, 0.40]; follow-up FDR-corrected paired t-test, corrected *p =* 0.01; effect size *d* = 0.55). Fig. 4b shows cross-patient decoding accuracies separated by target patient for a few sample patients (results for all target patients in Fig. S10). One patient (S3) in particular received an immense benefit from this additional cross-patient data (mean: aligned cross-patient = 0.53, patient-specific = 0.29), which we attribute to the initial lack of patient-specific data available (S3 had approximately one-third the amount of patient-specific data as the other patients), so the additional data provided critical repetitions to enable sufficient training. To validate that these increased cross-patient decoding accuracies were not due to the alignment procedure artificially inducing structure in cross-patient data that was not originally present, we generated surrogate data that preserved the first and second order statistical structure of our neural data (equal mean and covariance across trials, channels, and time) but was otherwise random (see Methods) and aligned this data to the target patient instead. We hypothesized that this surrogate neural data contained similar statistical structure but lacked speech-relevant latent dynamics and thus would fail to decode speech information from the target patient following alignment to the target patient’s space. Models trained on only this aligned surrogate data yielded accuracies with no significant difference from chance across all target patients (Fig. S11; Wilcoxon signed-rank test, W = 8, *n =* 8, *p* = 0.20), indicating alignment itself does not artificially yield improved phoneme decoding accuracy. These results show that speech decoding models can learn a shared representation of phoneme information from separate patient sources, substantiating the success of cross-patient speech BCIs.

**Figure 4.**
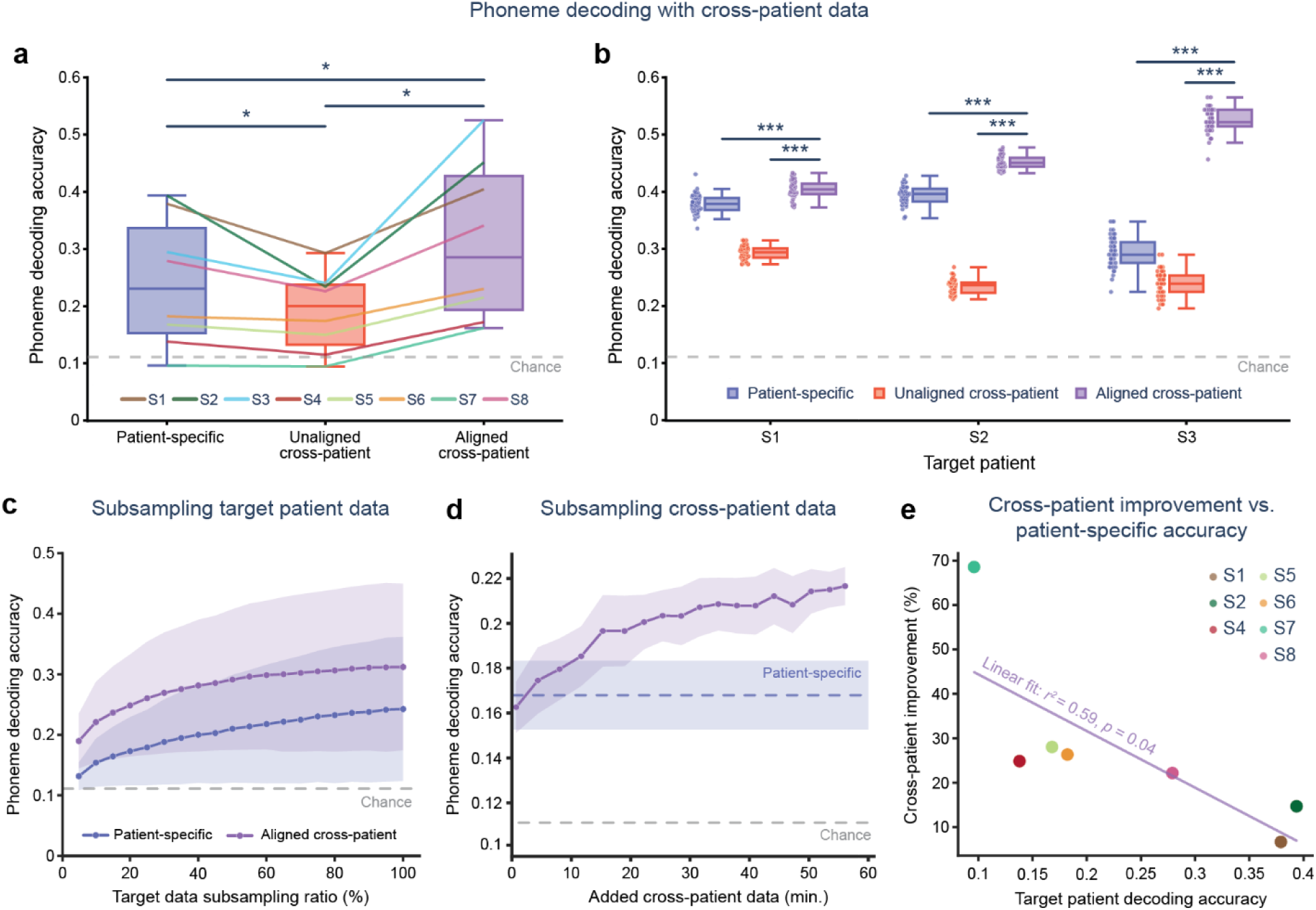
Expanded cross-patient datasets improve speech decoding accuracy. **a.** Cross-patient phoneme decoding across all eight patients. Each line represents cross-patient decoding aligned to a specific target patient’s space. In blue, SVM models are trained and tested on patient-specific data. In red, SVM models are trained on latent dynamics from all eight patients without alignment and tested on held-out patient-specific data from the target patient, resulting in a significant decrease in decoding accuracy (\**p < 0.05*, repeated measures ANOVA and follow-up FDR-corrected paired t-tests). In purple, SVM models are trained on latent dynamics from all patients with CCA alignment and tested on held-out patient specific data from the target patient, improving phoneme decoding accuracy relative to unaligned and patient-specific cases (*p < 0.05*, repeated measures ANOVA and follow-up FDR-corrected paired t-tests). The gray dashed line indicates chance levels of decoding. **b.** Cross-patient decoding for target patients S1, S2, and S3. Aligned cross-patient accuracies were significantly higher than both unaligned and patient-specific decoding contexts in each target patient (*\*\*\*p < 0.001*, post-hoc Tukey HSD test following one-way ANOVA). **c.** Patient-specific and aligned cross-patient decoding repeated as the percentage of data (trials) in the target patient dataset is subsampled. Accuracies shown here are aggregated across all patients (Fig. S16 for all patients separately). Even with limited target patient data to learn latent dynamics alignment, the cross-patient decoding outperformed patient-specific decoding (\*\**p < 0.01* at all subsampling ratios, exact two-sided permutation test). **d.** Aligned cross-patient decoding as the amount of additional cross-patient data (trials) is varied for patient S5. Little additional data was needed to increase the phoneme decoding accuracy beyond patient-specific levels. Diminishing gains in decoding accuracy are observed as additional cross-patient data is added. **e.** Relationship between cross-patient improvement (percent change between aligned cross-patient and patient-specific decoding) and target patient-decoding accuracy. Using a linear fit we found a significant negative relationship (*p <* 0.05, *r =* -0.77), showing cross-patient gain gets smaller as target patient-specific accuracy increases. Differences in aggregated results for all target patients were tested with a repeated measures ANOVA and follow-up paired t-tests with false discovery control. Within target patients, differences in distributions of phoneme decoding accuracy were compared across decoding contexts (patient-specific, unaligned cross-patient, aligned cross-patient) using a one-way analysis of variance (ANOVA) and post-hoc Tukey tests. Effect sizes were reported as Cohen’s *d.* For target patient data subsampling, an exact two-sided permutation test was performed at each target data subsampling ratio. FDR correction was applied across all subsampling ratio tests. A linear fit was performed between cross-patient gain and target patient-decoding accuracy to determine a significant relationship between the two variables.

When aligning the latent dynamics of one patient to those of a new target patient, it is unclear how much new data from the target patient must be collected to enable sufficient alignment to the target patient’s latent space. We expect that this alignment can be learned with minimal amounts of data – much less than needed to sufficiently train a complex patient-specific decoding model – as CCA is a linear alignment method. To test this, we repeated our cross-patient decoding analysis while subsampling the number of trials from the target patient at intervals of 5% (excluding S3 due to a previously mentioned lack of trials) used for alignment and decoder training (Fig. 4c, Fig. S16 for all patients separately). Across all patients, we found that CCA yielded successful alignment that enabled cross-patient speech decoding with significantly higher accuracies than patient-specific contexts as the subsampling ratio decreased (*n =* 8, *p =* 0.008 at all subsampling ratios with FDR correction, exact two-sided permutation tests), even for target patient datasets with only 5% of their original trials (∼7 trials/0.5 min.). This demonstrates the feasibility of cross-patient alignment with minimal data collection from new patients.

To quantify the relationship between the amount of additional cross-patient data and decoding performance, we repeatedly trained cross-patient SVMs while varying the amount of additional cross-patient data trials available. For each patient, we randomly subsampled a fixed number of trials equally from each other patient. We then aligned the subsampled trials to the target patient, trained a cross-patient SVM model, and evaluated its performance on a held-out test set from the target patient with 20-fold cross-validation. The results from this analysis for an illustrative patient (S5) are shown in Fig. 4d (all patients in Fig. S17). We chose to use S5 as the representative patient here, instead of S1 as we have done previously, as S5 better portrayed the increases in aligned decoding accuracy as additional cross-patient data was added that we saw across nearly every other patient. We found patient-specific performance (mean accuracy = 0.17) was exceeded after only 4.4 minutes of cross-patient data, with cross-patient decoding accuracy increasing to a mean of ∼0.22 (chance = 0.11) with all cross-patient data used. We observed diminishing gains in decoding accuracy as further cross-patient data was added for S5, as shown by a linear fit between the log-transformed amount of cross-patient data and aligned cross-patient decoding accuracy (*r^2^* = 0.94; *p =* 3.0*e-11*; Fig. S18). However, this may be an effect of the simplicity of the SVM models used (relative to state-of-the-art speech decoding models). We expect that more complex models would be able to better use this additional cross-patient data to further improve decoding accuracy.

We found varied improvements in aligned cross-patient decoding accuracy by patient. We hypothesized that this inter-patient variability may be related to the patient-specific decoding accuracy of the target patient, as patient-specific decoding models in patients with low accuracies may be insufficiently trained and stand to benefit more from additional cross-patient training samples. To investigate this, we plotted the relationship between each target patient’s patient-specific decoding accuracy and the percent change in decoding accuracy when trained with aligned cross-patient data relative to patient-specific data, which we denote the cross-patient improvement (Fig. 4e). For this analysis, we chose to exclude S3 due to the confound of dataset size, as it has one-third of the data of the other patients. Using a linear fit, we found a significant negative relationship between cross-patient improvement and patient-specific decoding accuracy of the target patient (*r* = -0.77, *p* = 0.04), indicating patients with lower patient-specific decoding accuracy saw a larger benefit to decoding accuracy with cross-patient data. We further investigated pairwise effects on inter-subject variability using a leave-one-out cross-patient training strategy (Fig. S19a; again, excluding S3). Specifically, for each target patient, we added all available cross-patient data to the target patient’s training set except for a single held-out patient and reported the percent decrease in cross-patient decoding accuracy relative to the accuracy with the full cross-patient training set. This allowed us to investigate changes in cross-patient accuracy on the level of a single source patient to look for additional drivers of inter-subject variability in cross-patient accuracy. Using linear regression, we found a significant relationship between the patient-specific decoding accuracy of the target patient and the left-out change in cross-patient accuracy (Fig. S19b; *r^2^* = 0.51, FDR-corrected *p* = 3.89*e*-7), indicating that target patients with higher patient-specific accuracy are more robust to left-out data than target patients with lower patient-specific accuracy. Surprisingly, we found no significant effect of left-out patient-specific accuracy (Fig. S19c; *r^2^* = 0.05, FDR-corrected *p* = 0.24) or distance between electrode arrays in neuroanatomical space (Fig. S19d; *r^2^* = 0.01, FDR-corrected *p* = 0.64; S4 and S7 additionally excluded to constrain analysis to left hemisphere). These results align with Fig. 4e and the view that insufficient training of these lower accuracy models results in more benefit from (or more reliance on) cross-patient data.

### Cross-patient alignment improves speech decoding in a simulated real-time environment

Thus far, the presented alignment and decoding analyses were performed in offline decoding contexts that use non-causal windows of neural activity centered around patient responses. However, speech BCI systems must operate causally and without access to patient response timing. As such, the translatability of our cross-patient alignment methodology to more relevant real-time speech decoding contexts remains unclear.

To investigate the effects of cross-patient alignment on real-time speech decoding performance, we applied an adapted alignment and decoding strategy on simulated real-time neural recordings from our intraoperative dataset. We first extracted HG power within short windows of the neural recording (20 ms, as in previous real-time speech BCI work^7^) for each trial. HG data from a designated training period was used to learn decomposition and alignment matrices between source and target patients with our PCA-CCA alignment method, which we used to increase the size of the target patient’s training dataset with aligned data from source patients as in Fig. 4. We then trained a recurrent neural network (RNN) model with connectionist temporal classification^35^ (CTC) loss to output a timeseries of phoneme predictions from causal, sliding windows of HG power without knowledge of patient speech onset timing (Fig. 5a). To evaluate performance of the CTC-RNN model with and without cross-patient alignment, we analyzed the phoneme error rate (PER) between predicted and true phoneme sequences on a held-out test set from the target patient when the model was trained on patient-specific, unaligned cross-patient, and aligned cross-patient data.

**Figure 5.**
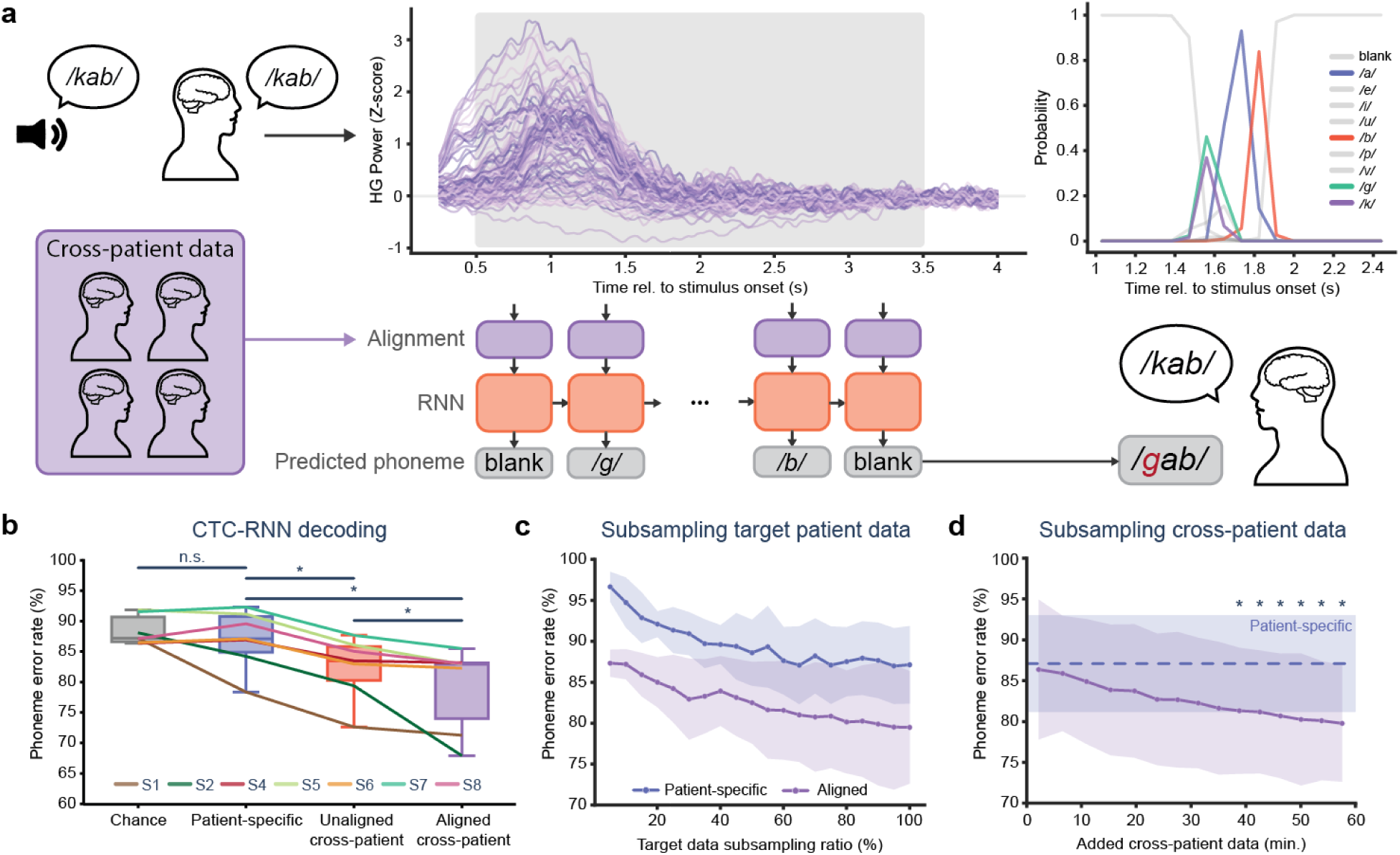
Cross-patient alignment improves real-time speech decoding performance. **a.** Schematic of simulated real-time speech decoding pipeline. (Left) High-gamma (HG) signals are extracted in 20 ms windows to simulate streaming of real-time features. We use a sliding, causal decoding window (280 ms, 80 ms stride) to predict a timeseries of phonemes from 0.5 to 3.5 seconds following stimulus onset. A decoding model (recurrent neural network) is trained with a combination of patient-specific and aligned cross-patient data. We use connectionist temporal classification (CTC) loss to train the model to predict a sequence of phonemes from the output timesteps. (Right) Phoneme predictions from a sample trial. The top plot shows the timeseries of phoneme output probabilities. The trial starts with blank predictions during silence, then predicts a sequence of three phonemes (*/g/, /a/, /b/*) after. The bottom diagram shows the predicted sequence (*/gab/*) compared to the true sequence (*/kab/*). The mistakenly predicted first phoneme results in a phoneme error rate (PER) of 33%. Despite this mistake, the predicted probability of the */k/* phoneme is nearly that of the */g/* phoneme, indicating confusion between these similar phonemes. **b.** CTC-RNN decoding performance, measuring phoneme error rate between predicted and true sequences. There is no significant difference between chance and patient-specific PERs (*p = 0.81*, FDR-corrected Wilcoxon signed-rank test). However, unaligned cross-patient PERs are significantly lower than patient-specific (\**p < 0.05*, FDR-corrected Wilcoxon signed-rank test) and aligned cross-patient PERs are significantly lower than both patient-specific and unaligned PERs (*p < 0.05*, FDR-corrected Wilcoxon signed-rank test. **c.** PER in patient-specific and aligned cross-patient contexts as the percentage of target patient data (trials) is subsampled, averaged across all patients. Cross-patient PERs are significantly lower than patient-specific PERs for all subsampling ratios (*p < 0.05*, FDR-corrected Wilcoxon signed-rank test). Cross-patient gains remain even with as little as 5% of the original patient-specific dataset. **d.** PER in aligned cross-patient decoding as the amount of additional cross-patient data is varied, averaged across all patients. PER starts at patient-specific values with little cross-patient data, then steadily decreases as more cross-patient data is added. Aligned cross-patient PERs are significantly lower than patient-specific PERs after ∼38.8 minutes of cross-patient data are added (*p < 0.05*, FDR-corrected Wilcoxon signed-rank test). Differences between chance, patient-specific, unaligned, and aligned CTC-RNN decoding was performed with multiple FDR-corrected Wilcoxon signed-rank tests. At each 5% interval in the target patient subsampling analysis, a Wilcoxon signed-rank test was performed to assess differences in error rates between aligned and patient-specific results across all patients. FDR-correction was performed across all subsampling points. Similarly, for the cross-patient subsampling analysis, Wilcoxon-signed rank tests were performed at each added cross-patient data value to compare aligned cross-patient performance to the single distribution of patient-specific decoding performance. FDR-correction was again performed across all subsampling points.

We again found that decoding models trained on aligned cross-patient data outperformed both those trained on patient-specific and unaligned cross-patient data (Fig. 5b, all target patients separately in Fig. S20). Across all patients (excluding S3 due to a limited data amount – see Methods), we found no significant difference between PER in a chance distribution, obtained from shuffling label identity across trials, and patient-specific training (mean: chance = 88.4%, patient-specific = 87.1%; Wilcoxon signed-rank test, *W* = 12, *n =* 7, FDR-corrected *p* = 0.81), likely due to our limited amount of data paired with the complex CTC-RNN decoding model resulting in insufficient training. Surprisingly, we found that CTC-RNNs trained on unaligned cross-patient data achieved significantly lower PERs than in the patient-specific context (mean unaligned cross-patient = 82.5%; Wilcoxon signed-rank test, *W* = 0, *n =* 7, FDR-corrected *p* = 0.02). This was the opposite of the effect that we observed in our offline decoding analysis using SVM models (Fig. 4a), suggesting feature transformations within the RNN model may better elucidate shared patterns across data from unaligned patient sources than simpler SVM models. With models trained on aligned cross-patient data, we saw a further decrease in PER relative to the unaligned context across all patients (mean aligned cross-patient = 79.4%; Wilcoxon signed-rank test, *W* = 0, *n =* 7, FDR-corrected *p* = 0.02). We saw particularly strong improvements when using aligned cross-patient data in S2, with PER decreasing from 84.2% in the patient-specific context to 67.9% in the aligned cross-patient context. These results confirm that our observed increases in speech decoding performance following cross-patient alignment translate to real-time decoding contexts.

To quantify changes in CTC-RNN performance with variable amounts of data, we subsampled target and cross-patient data while repeating our simulated real-time alignment and decoding method. First, we subsampled the amount of target patient data available to CTC-RNN models trained on either patient-specific or aligned cross-patient data at intervals of 5% (Fig. 5c, all patients separately in Fig. S21). We found that aligned cross-patient decoding significantly outperformed patient-specific decoding for all target data subsampling ratios (Wilcoxon signed-rank test, *n* = 7, *W* = 1, FDR-corrected *p* = 0.03 for a subsampling ratio of 60%, *W* = 0, FDR-corrected *p* = 0.02 for all other subsampling ratios). Importantly, even with only 5% of the original patient-specific data (∼5 trials/0.3 min.), aligned cross-patient decoding outperformed patient-specific decoding. As such, there is little “barrier to entry” required in the data collection before cross-patient gains are observed. Next, given a full patient-specific dataset (∼6.3 min. experiment duration – see Methods), we evaluated decoding performance as we subsampled the amount of cross-patient data added to test the necessary amount of cross-patient data needed to outperform patient-specific decoding (Fig. 5d, all patients separately in Fig. S22). We found that aligned cross-patient decoding achieved significantly lower PERs than patient-specific decoding after ∼38.8 minutes of cross-patient data were added (Wilcoxon signed-rank test, *n* = 7, *W* = 0, FDR-corrected *p* = 0.04 for cross-patient data > 38.8 min., *p* > 0.05 for lesser amounts of cross-patient data). To estimate performance as additional cross-patient data is added, we performed a linear fit between the amount of added cross-patient data and log-transformed PER (*r^2^* = 0.98, *p* = 6.6*e-12,* Fig. S23). Using this fit, we would achieve a 25% average PER with 14.6 hours of cross-patient data and a 10% average PER with 25.3 hours of cross-patient data. These results align well with learning curves composed entirely of patient-specific data shown in an ECoG speech BCI in previous literature by Metzger and colleagues^8^, suggesting that cross-patient data can be a valid substitute for long-term collection of patient-specific data.

To ensure the timing required to learn cross-patient alignment transformations is feasible in real-time contexts, we quantified the time to learn CCA transformation matrices for each target patient. We found an average time of 5.8 seconds, with up to ∼10 seconds maximum, to calculate alignment between a target patient with dataset size of ∼6.3 min. and ∼60 min. of cross-patient data (Fig. S24a). This alignment only needs to be learned once following the collection of training data for the target patient. As such, this time should be framed in the context of the overall experiment duration, in which a one-time delay of 10 seconds is negligible compared to the 6.3 min. of prior data collection, meaning that learning CCA alignment can easily be integrated into data collection paradigms. Further, we have taken the approach of aligning cross-patient data to the space of the target patient, resulting in no change to real-time prediction latency, as there is no need to transform the real-time-acquired data from the target patient during evaluation periods prior to input to the decoding model. However, to account for cases in which the user may wish to align to a space other than the target patient, thus necessitating a linear transformation by a learned alignment matrix, we quantified the additional latency required for a linear transformation of the input data prior to input to the CTC-RNN (Fig. S24b). We found an average timing of 0.04 ms to apply a linear transformation to the acquired target patient data. In comparison to the CTC-RNN prediction latencies ranging from 1 to 7 ms this was a negligible increase to prediction latency, especially given the 80 ms latency budget defined by the stride of our decoding window. Overall, timing requirements of cross-patient alignment via CCA are compatible with requirements of real-time data collection and latency budgets.

### High-resolution and broad coverage are critical to alignment

To fully assess the benefits of high spatial sampling and coverage for cross-patient decoding, we quantified the spatial sampling of the SMC required to sufficiently elucidate speech-relevant latent dynamics that can be aligned across patients. We subsampled various spatial characteristics of our μECoG arrays and calculated both the cross-patient decoding accuracy of SVM models trained on aligned latent dynamics and the cross-patient representational similarity (via RSA) in the extracted latent dynamics at each subsampled point. We separately investigated the influence of subsampling electrode density, array coverage, and electrode contact size.

To investigate the effects of electrode density, as represented by the distance between electrodes (pitch), we used Poisson disk sampling to randomly sample individual electrodes across the array while constraining the minimum distance between sampled electrodes to approximate the cortical sampling of lower-density ECoG arrays. Simulated pitch ranged from 10 mm, the pitch of standard clinical ECoG devices, to 1.5 mm. To quantify the speech information encoded in the pitch-subsampled data, we repeated our cross-patient phoneme decoding analysis from Fig. 4 at each effective pitch value. We observed increases in phoneme decoding accuracy as pitch decreased (an increase in spatial sampling) in both patient-specific and aligned cross-patient contexts (Fig. 6a, top; Fig. S25 for all patients separately). Crucially, we found that aligned cross-patient models significantly outperformed patient-specific models for only pitch values under 3 mm (*n =* 8, FDR-corrected *p =* [0.04, 0.04] for pitch ɛ [1.5, 2] mm, respectively, *p > 0.05* for all other pitch values, exact two-sided paired permutation test). We additionally found that the representational similarity between aligned latent dynamics of separate patients increased as the effective pitch decreased across all patients, with more rapid increases for smaller pitch values (Fig. 6a, bottom). These results indicate that sufficient sampling of cross-patient latent dynamics necessitates high-density neural interfaces.

**Figure 6.**
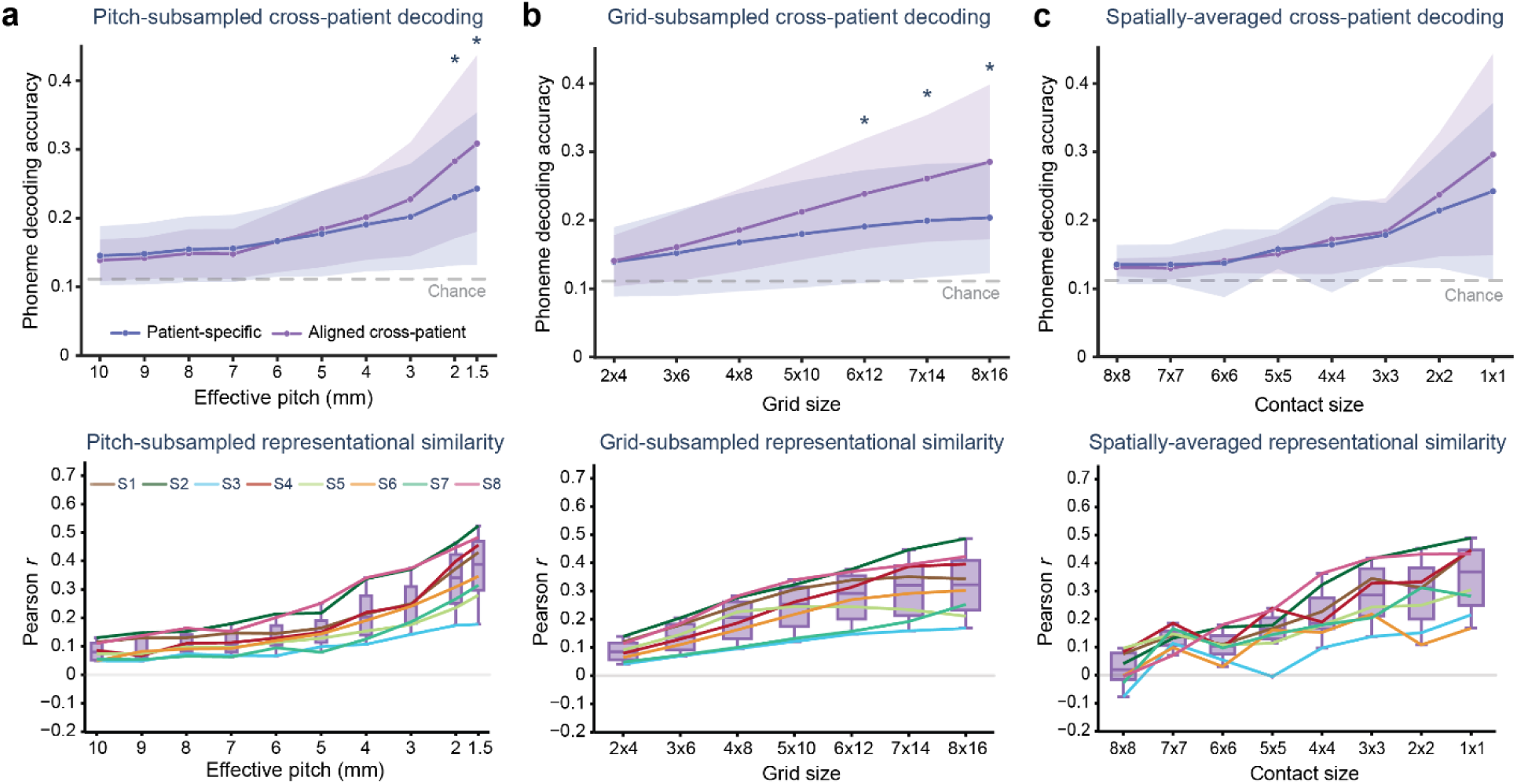
High-resolution sampling with broad coverage is required for successful cross-patient alignment. **a.** Effect of electrode density. (Top) Cross-patient phoneme decoding with varied pitch (distance between electrodes). Accuracy increases across patient-specific and aligned cross-patient decoding contexts as pitch decreases. Aligned cross-patient accuracies were only significantly higher than patient-specific accuracies for effective pitch less than 3 mm (*p < 0.05* for pitch ɛ [2, 1.5] mm, exact two-sided paired permutation test, *p*<0.05*). (Bottom) Cross-patient representational similarity, as measured by the Pearson correlation between representational dissimilarity matrices, increases as the pitch decreases. Each line represents a single target patient, where correlations are averaged across all non-target patients aligned to the target patient. **b.** Effect of array coverage. As in **a,** but for subsampled grids of varying size. (Top) Phoneme decoding accuracy similarly increases as grid size increases. Aligned cross-patient accuracies were only significantly higher than patient-specific accuracies for grid sizes greater than 6x12 (∼8 mm x 17 mm*; p < 0.05* for grid sizes ɛ [6x12, 7x14, 8x16], exact two-sided paired permutation test). (Bottom) Cross-patient representational similarity increases as grid size increases. **c.** Effect of contact size. As in **a,** but with varying contact sizes obtained by spatially-averaging different amount of neighboring micro-contacts. (Top) Phoneme decoding accuracy increases as contact size decreases. The mean phoneme decoding accuracy of aligned cross-patient models is higher than patient-specific models for contact sizes under 3x3 (∼3 mm diameter) however there is no significant difference after correction for multiple comparisons (*p > 0.05* for all contact sizes, exact two-sided paired permutation test). (Bottom) Cross-patient representational similarity generally increases as contact size decreases. Statistical testing was performed to compare differences between aligned cross-patient and patient-specific decoding accuracies for different subsampling methods. For each subsampling method (pitch, grid, and spatial-averaging) we performed an exact two-sided, paired permutation test at each subsampled point with FDR correction across all points.

To evaluate the necessity of neural interfaces with broad coverage of the SMC, we selected sub-grids of electrodes from our full electrode arrays and repeated the analyses in Fig. 5a. Grid sizes ranged from 2x4 (∼2 mm x 5 mm) to 8x16 electrodes (∼11 mm x 23 mm). Grid-subsampled phoneme decoding gradually increased as coverage increased for both decoding contexts (Fig. 6b, top; Fig. S26 for all patients separately). Aligned cross-patient models had significantly higher accuracies than patient-specific contexts for only grid sizes 6x12 and larger (*n =* 8, FDR-corrected *p =* [0.04, 0.04, 0.04] for grid sizes ɛ [6x12, 7x14, 8x16], respectively, *p > 0.05* for all other grid sizes, exact two-sided paired permutation test), which corresponds to coverages greater than ∼8 mm x 17 mm, showing that broad coverage of the SMC is critical for successful sampling of cross-patient latent dynamics. Similarly, we found that the representational similarity between aligned cross-patient latent dynamics increased as the size of the subsampled grids increased across all patients, showing better similarity with broader coverage (Fig. 6b, bottom).

Finally, we quantified the effects of contact size by extracting HG information from recorded signals averaged together across neighboring electrode contacts to approximate larger contacts. Contact sizes ranged from 8x8 micro-contacts (∼11 mm diameter) to a single micro-contact (200 μm). We found that decoding accuracy increases as contact size decreases for both decoding contexts (Fig. 6c, top; Fig. S27 for all patients separately). While the decoding accuracy of cross-patient SVMs was higher than the patient-specific decoding accuracy for contact sizes under 3x3 (∼3 mm), there was no significant difference after correction for false discovery rate (*n =* 8*, p > 0.05* for all contact sizes, exact two-sided paired permutation test). Representational similarity in aligned latent dynamics generally increased as contact size decreased (Fig. 6c, bottom). These results suggest that electrode density and array coverage play a more important role in properly sampling SMC cross-patient latent dynamics during speech.

## Discussion

Speech BCIs have recently demonstrated impressive performance in rapidly and accurately translating neural signals to words in patients with an inability to speak due to neuromotor disorders. However, these BCIs typically require long periods of patient-specific data collection that limit their deployability and effectiveness. In this work, we investigated shared, cross-patient representations of speech production in the SMC to reduce the deployment time of speech BCIs by enabling cross-patient decoding models. While cross-patient speech decoding has been previously investigated using transfer learning^14^, as of this writing, this is the first study, to our knowledge, to use neural activity aligned across human patients for speech decoding. Our results support the idea that cross-patient models can reach sufficient data requirements for advanced decoding methodologies with minimal data collection from any individual patient and can be easily transferred across patients, resulting in robust and rapidly deployable speech BCIs that are more accessible to a broader range of patients.

A novel aspect of this study is the investigation of shared latent dynamics in human speech. Previously, the alignment of latent dynamics across individuals has been constrained to non-human primates and rodents performing limb movement tasks^26^. Here, we show this phenomenon of shared neural activity across individuals extends to the human motor speech system, where the implications for translational BCI contexts stand to provide substantial benefits for training machine learning models and accelerating deployment following device implantation. Despite differences between limb movement and speech production, the presence of shared speech latent dynamics is a logical extension as both movement contexts pertain to the alignment of low-level neural motor representations underlying highly-coordinated muscle movements.

We hypothesized that neural population activity underlying speech articulator movement is well-represented in patients’ latent dynamics and shared across patients. This is an important motivation for the use of functional alignment in our study, as the transformations learned through our alignment procedure do not simply reflect efforts of domain adaptation between a target and source patient. Instead, this method aims to exploit the existence of a shared neural manifold of speech motor control to account for differences in neural sampling across patients that arises from variability in neuroanatomy and electrode placement. This distinction is critical, as it allows for principled pooling of neural data across patients while maintaining a consistent decoding model, and provides a complementary alternative to existing transfer learning strategies in neural decoding.

To extract latent dynamics from HG activity across many channels, we used PCA as a dimensionality reduction method to approximate latent dynamics. Further, to perform alignment between latent dynamics from different patients, we used CCA to learn linear transformations between latent dynamics. We chose these methods for their simple and interpretable transformations, where aligned latent dynamics are the result of spatial filters applied to the original electrode space. However, PCA and CCA are limited because they 1) fail to capture relevant nonlinear latent representations of cortical neural activity during motor control^24,39^ 2) treat data from different time points as independent observations, which discards potential temporal information that could be useful in learning a better alignment across patients. Recent dynamical systems frameworks have been developed that use deep representation learning to elucidate nonlinear latent dynamics while explicitly modeling temporal information, such as Latent Factor Analysis for Dynamical Systems (LFADS)^39^ and MAnifold Representation Basis LEarning (MARBLE)^40^. Future studies should investigate nonlinear cross-patient representations of speech production extracted from these advanced dynamical systems models to determine if these changes improve alignment quality between patients. However, study designs must be wary of limited interpretability, relative to linear decomposition methods, with deep learning models. Additionally, the amount of data per patient required to successfully learn latent dynamics should be considered in these deep models, as a steep data requirement to learn and align latent dynamics is antithetical to the desired rapid deployability for translational BCI contexts.

We found that alignment of latent dynamics by CCA preserved spatial properties of speech information. In many cases, we found strong correlations between ground-truth articulator maps and articulator maps projected to that patient’s space from another source patient’s space. However, we found some cases of negative correlation, and thus misalignment, when reconstructing articulator maps with weak representations, suggesting this alignment may fail to properly capture weakly represented speech information. Statistically, CCA alignment is designed to maximize correlation between cross-patient latent dynamics but does not enforce a one-to-one spatial correspondence. As a result, reconstruction into an individual patient’s electrode space can introduce distortions, particularly when the shared latent space captures only a subset of the variance present in the target patient. As described earlier, this may be a failure of the linear latent dynamics being unable to approximate necessary nonlinear activity to accurately preserve observed spatial activations^24^. From a neurophysiological perspective, these discrepancies may reflect true, minor inter-patient variability in speech motor representations. While similar cortical regions are engaged across individuals, the combined differences in anatomy, electrode coverage, and functional organization may not be fully explained by a shared motor control manifold. As such, these observations suggest that while a shared latent structure supports cross-patient generalization, it may not fully capture all patient-specific features of neural organization.

Regardless, we found that this alignment enabled successful cross-patient decoding, with models trained on additional cross-patient data consistently outperforming those trained on patient-specific data alone. In some patients, these improvements were substantial. These analyses, however, were conducted on a relatively small dataset, combining recordings from eight patients with limited data per individual. We expect that scaling to larger and more diverse patient cohorts will further enhance decoding performance, particularly when paired with more advanced models such as recurrent neural networks (RNNs) and other deep learning approaches.

By systematically varying the composition of the cross-patient training set, we observed significant inter-patient variability in decoding gains. Notably, the strongest predictor of improvement was the baseline decoding accuracy of the target patient, whereas neither the identity of source patients nor the neuroanatomical distance between electrode arrays had a significant effect. However, this neuroanatomical distance analysis did not consider any effect of cortical regions-of-interest (ROIs), which we suspect play an important role in finding shared or distinct latent dynamical representations of speech. It would be difficult to implement such an analysis with this dataset due to the homogeneity of motor area coverage with our patients’ arrays. However, we believe future work should consider a comprehensive ROI-based analysis with stereo-electroencephalography (SEEG), as the distributed coverage of SEEG recordings would be well suited to answer this question, assuming electrode density and sampling within ROIs are sufficient to extract relevant speech-related latent dynamics. Overall, these findings suggest that cross-patient alignment provides the greatest benefit when patient-specific models are limited by insufficient data or lower signal quality, effectively acting as a form of data augmentation that stabilizes decoding in these settings.

Our CTC-RNN results demonstrate that cross-patient alignment can substantially improve real-time speech decoding performance in contexts that more closely reflect operational BCI systems. Using simulated real-time μECoG recordings and causal sliding windows of HG activity, we found that recurrent neural network models with CTC loss trained on aligned cross-patient data consistently outperformed both patient-specific and unaligned cross-patient models. Importantly, these benefits were observed even with minimal amounts of target patient data, indicating that cross-patient alignment can reduce the data collection burden required to achieve accurate decoding. Performance improvements scaled predictably with the amount of aligned cross-patient data, and log-extrapolation to long-term dataset sizes suggested that cross-patient data could serve as a viable substitute for extensive patient-specific training.

Critically, the latency required to compute alignment transformations and apply them to incoming neural data is minimal relative to the overall experimental timescale and decoding window, demonstrating that this approach is compatible with real-time BCI constraints. These findings suggest that cross-patient alignment provides a practical and effective means to leverage shared neural structure across individuals, stabilizing decoding in low-data regimes while maintaining real-time feasibility. Together, these results underscore the potential of cross-patient approaches to accelerate BCI deployment, reduce patient-specific training requirements, and support robust speech decoding in clinically relevant contexts.

By subsampling various spatial characteristics of our μECoG arrays, we found that neural interfaces with both high resolution and broad coverage of the SMC are best suited to align latent dynamics and train cross-patient speech decoding models. Neural interfaces in current speech BCIs^7,8^ typically prioritize only one of these characteristics, as intracortical microelectrode arrays offer high electrode density but limited coverage and ECoG grids offer wider coverage but with limited spatial sampling. While our results suggested potentially limited generalization to non-μECoG neural interfaces because of degraded sampling of the neural signal, previous work by Safaie and colleagues has shown successful alignment of motor latent dynamics across multiple individuals from neural recordings with small intracortical microelectrode arrays^26^, showing feasibility with reduced coverage. Additionally, future studies should consider alignment across neural interface types. As cross-patient alignment is only calculated on dimensionality-reduced latent dynamics, alignment could still be successful with a heterogeneity of neural interfaces across patients, which would increase data availability for cross-patient alignment. Nonetheless, our current findings motivate the use of μECoG for cross-patient speech decoding as it enables both dense and broad sampling.

It is important to note that the current study is limited to the analysis of cross-patient latent dynamics during overt speech in able-bodied speakers. These patients differ functionally from individuals with impaired speech function due to neuromotor disorders like ALS or brainstem stroke, where neural signal related to speech may be weaker, due to potential degradation of cortical structures necessary for speech production and monitoring^41,42^, or organized differently than those with intact speech as a whole due to compensatory reorganization of neural motor pathways^43,44^. As such, it remains unknown if the preservation of speech latent neural dynamics that we observed here (in individuals without speech dysfunction) will be present across individuals with varying levels of motor system impairments. However, recent speech BCI studies have found that articulatory representations of speech may be well preserved in patients with impaired speech many years after diagnosis or injury with ALS^7^ or brainstem stroke^8^, respectively, suggesting that the shared articulatory latent dynamics we observed here may indeed be relevant to the translational development of speech BCIs.

Together, our results demonstrate that speech-relevant neural population dynamics are shared across human patients, providing a foundation for more efficient and scalable speech BCI systems. With cross-patient alignment based on latent dynamics, future speech BCIs may achieve faster adaptation with minimal patient-specific data, resulting in improved patient usability and performance. Most importantly, cross-patient speech BCIs will improve communicative ability and quality of life for individuals with paralysis from neuromotor disorders.

## Methods

All our research studies were approved by the Institutional Review Board of the Duke University Health System under the following protocol ID: Pro00072892: Studying human cognition and neurological disorders using µECoG electrodes.

### Participants

This study included neural data collected intraoperatively from eight awake patients (mean age = 57.4, one female) with fully intact speech at the Duke University Medical Center (DUMC). Intraoperative recordings were performed while patients S1, S2, S4, and S5 underwent surgical interventions for movement disorders by deep brain stimulator (DBS) implantation and patients S3, S6, S7, and S8 were undergoing brain tumor resection. All subjects were native English speakers. Written informed consent was obtained prior to surgery in accordance with the Institutional Review Board of the Duke University Health System.

### Intraoperative neural recording

Intraoperative neural recordings were performed using custom-designed micro-electrocorticographic (μECoG) arrays that were fabricated by Dyconex Micro Systems Technology (Basserdorf, Switzerland). Recordings were performed during awake neurosurgical procedures for movement disorders (deep brain stimulator implantation – DBS) and tumor resection. Two different μECoG designs were used. First, for DBS implantation surgeries, a narrow (11 mm x 21 mm) μECoG array with 128 micro-contacts (200 μm contact diameter, 1.33 mm center-to-center spacing between contacts) was inserted through a burr hole and placed in subdural space over the target sensorimotor cortex (SMC). For tumor resection surgeries, larger (38 mm x 21 mm) μECoG arrays with 256 micro-contacts (200 μm contact diameter, 1.72 mm center-to-center spacing between contacts) were placed subdurally, directly over the SMC through a craniotomy window. Both array types used gold electrodes, with a platinum-iridium coating (in all patients except S1), embedded in a liquid crystal polymer thin-film (LCP-TF) substrate (Impedance information for all arrays in Fig. S2). Arrays were sterilized with ethylene oxide gas prior to surgical implantation. μECoG arrays were connected to custom headstages using Intan Technologies RHD chips to amplify and digitize recorded neural signals. These headstages were then connected to an Intan Technologies RHD recording controller via micro high-definition multimedia interface (μHDMI) cables. Recorded neural data were processed by the Intan RHX v3.0 data acquisition software (Intan Technologies, Inc.). Recordings were analog filtered between either 0.1 Hz and 7,500 Hz or 0.1 Hz and 10,000 Hz. Recordings were digitized at 20 ksps.

To visualize μECoG array location for each patient, electrode locations were co-registered with patient cortical reconstructions that were generated from preoperative MRI images using Freesurfer’s *recon-all-clinical* function^45^. For DBS implantation surgeries, array location was confirmed with an intraoperative CT scan following recording that was aligned to the reconstructed preoperative MRI volume. Electrodes were localized from the CT scan using BioImage Suite^46^. For tumor resection surgeries, array location was registered to the preoperative MRI with Brainlab Neuronavigation (Munich, Germany) coordinates marked at array corners. Brainlab coordinates were imported into BioImage Suite, where intermediary electrode locations were interpolated to match the known electrode spacing of the electrode array (due to missing Brainlab coordinates for subject S8, electrode location was manually determined based on intraoperative images pre- and post-array placement). In both DBS and tumor resection cases, electrode locations were subsequently projected to the leptomeningeal surface of the brain. Electrode arrays of all patients were visualized together by projection to a common brain (MNI-152, Fig. 1) and separately on patient-specific brains (Fig. S1).

### Speech task

Patients performed a speech repetition task in the acute intraoperative neural recording setting. In a single trial, patients are presented with an auditory stimulus representing one of 52 unique non-words via a computer speaker and asked to verbally repeat the sounds they hear. Non-words were sequences of three phonemes designed to sample the phonotactics of American English. Sequences were composed of three of nine possible phonemes (four vowels: */a/, /e/, /i/, /u/* and five consonants: */b/, /p/, /v/, /g/, /k/*) arranged in consonant-vowel-consonant (e.g. */k/, /a/, /b/*) or vowel-consonant-vowel (e.g. */a/, /b/, /e/)* format (See Table S2 for all stimuli). Trials were composed of stimulus presentation (stimulus duration: ∼0.5 s) followed by a three second response window with 250 ms jitter in between trials. Patients responded ∼1.1 s after the beginning of the stimulus presentation, with their response lasting ∼0.45 s. Trials were presented across three consecutive blocks (52 trials per block for each unique non-word token). Each block corresponded to ∼0.5 min. of utterance duration. Stimuli order was randomly shuffled within blocks. Trial start times were marked using flashing lights recorded by a photodiode on the task presentation laptop (Thorlabs, Inc.). Patient responses were recorded with a lavalier microphone (Movophoto, Inc.) connected to a pre-amplifier (Behringer). Both photodiode and microphone data were sent via a BNC cable/connector to auxiliary channels on the Intan RHD recording controller for synchronization with recorded neural data at 20 ksps. The speech task was designed using Psychtoolbox in MATLAB R2014a. Phoneme labels (identity, start time, stop time) for both auditory stimuli and patients responses were first automatically generated using the Montreal Forced Aligner^47^ and then manually corrected using the Audacity audio-editing software^48^.

### Neural data preprocessing

Recorded neural data was decimated to 2 ksps using a finite impulse response anti-aliasing filter (Kaiser window, MATLAB’s *resample* function) and common-averaged-referenced (CAR) to remove common noise across channels. Electrodes with large impedance values (>1 Mohm) were excluded from CAR. After CAR, trials with large transient activity (>50 μV between timepoints) were discarded. All preprocessing was done in MATLAB R2021a.

### Spectrogram visualization

Spectrograms were calculated for each electrode to visualize spectrotemporal neural activity during patient responses (Fig. 1d-e). Spectral information was extracted using 500 ms windows with a 50 ms step-size and 10 Hz frequency smoothing using a multi-taper time-frequency analysis method^49^. Spectrograms during patient responses were normalized by a baseline period 500 ms prior to auditory stimulus presentation by dividing response spectrograms by the average baseline power-spectral density to remove *1/f* broadband activity. Normalized response spectrograms were averaged across trials for each electrode and the log-power was calculated around patient response onset (-1 to 2 s) to yield neural activity in units of decibels.

### High-gamma extraction

The high-gamma band (HG: 70-150 Hz) of the neural signal was extracted to estimate neural population dynamics in the SMC. HG signals have previously been shown to encode rich speech production information^8,19,31^ and strong correlations with neuron spiking activity^32–34^, suggesting a valid estimation of speech-relevant SMC population dynamics. Following preprocessing, neural signals were bandpass filtered using frequency-domain Gaussian filters centered at eight logarithmically spaced center frequencies (with logarithmically scaled bandwidths around each center frequency) between 70 and 150 Hz^50^. Specifically, the neural timeseries were decomposed into their corresponding frequency representations with the fast Fourier transform (FFT) and multiplied in frequency space with a Gaussian kernel centered around each center frequency to select and smooth information from the frequencies in the desired band. The inverse FFT was then performed to convert bandpass filtered signals at each band back to the time domain. Envelopes of signals in each band were calculated as the absolute value of the Hilbert transform and subsequently averaged across the eight bands. The resultant timeseries was then downsampled to 200 Hz with a finite impulse response anti-aliasing filter (Kaiser window, MATLAB’s *resample* function). For visualization of HG activity across arrays (Fig. 2a), HG activity was z-score normalized to the baseline period (500 ms prior to auditory stimulus presentation) for each trial. For all other analyses (latent dynamics extraction, decoding), HG activity was normalized by subtracting the average HG envelope of the baseline period. Analyses in this work used HG activity in a one second window (-500 ms to 500 ms) around patient response onset. Channels with significant speech HG activity were determined by statistical comparison between response and baseline periods (see *Statistical Analysis*). HG extraction was performed in MATLAB R2021a.

### Generation of surrogate data

As a control condition for many analyses, we repeated these analyses using data generated from the Tensor Maximum Entropy (TME) procedure^51^ to for each patient instead of their true data. In the following text, we briefly adapt the description of the method as presented by Elsayed and Cunningham. TME-generated surrogate data preserves the second-order statistics (mean and covariance) of the data with respect to each dimension of the data (trials, time, and channels), while otherwise being maximally random. To perform this, the method generates a probability distribution *p* that has maximum entropy *H* under the constraints of equal mean and variance to the original data for each dimension. That is, for surrogate data *D_surr_* composed of *s* trials, *t* timepoints, and *n* channels (shape *=* (*s, t, n*)), we have:

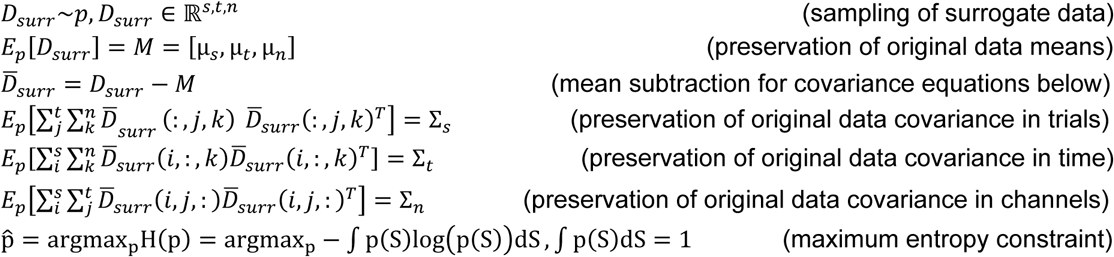

Where μ_x_ and Σ_x_ are the mean and covariance for dimension *x* of the data (trials, time, channels), respectively, 𝐸_p_[.] is the expectation over probability distribution *p,* and *p^* is the optimal probability distribution to sample surrogate data, which is essentially a 3D Gaussian distribution with standard deviation controlled by the covariance preservation and maximum entropy constraints.

As these constraints only impose structure with respect to covariance in each dimension, and randomness elsewhere, latent dynamics extracted from TME data should not encode speech-relevant information. As such, our control analyses allow us to evaluate alignment without the effect of speech information while still controlling for important aspects of the neural signal such as autocorrelation effects and correlation between channels. Surrogate data generation was performed with code from https://github.com/gamaleldin/TME in MATLAB R2021a.

### Latent dynamics extraction

We extracted latent dynamical activity from estimated SMC population activity as measured in HG activity around response onset for each electrode and trial (data array of the dimensions – *s, t, n*), where *s* is the number of trials, *t* is the number of timepoints, and *n* is the number of significant electrodes/channels). We used principal component analysis (PCA, scikit-learn^52^) to reduce the high-dimensional channel space into a low-dimensional space composed of *k* principal components (PCs), where each PC represented a linear combination of activity across the *n* channels. *k* was determined as the number of components explaining 90% of the variance in the original data. This yielded patient-specific latent dynamics (data array of the dimensions – *s*, *t, k*) as the projection of the original HG activity on the *k*-dimensional latent space. Extraction of latent dynamics was performed in Python 3.10.

Clustering of articulatory features in the latent space was visualized by further projecting latent dynamics into a two-dimensional subspace (Fig. 2b) using t-distributed Stochastic Neighbor Embedding (t-SNE, scikit-learn, perplexity = 30), a widely used technique for visualization of high-dimensional data^53^. t-SNE decomposition of latent dynamics yielded a data array of dimensions (*s*, 2). Visualization was performed by plotting each trial as a point in 2D t-SNE space and coloring each trial point by its articulatory category. Quantification of clustering strength was performed by calculating the Silhouette Coefficient (scikit-learn, distance = Euclidean) for each trial in t-SNE space, which measures the difference between inter- and intra-cluster distances^54^. The score is more positive when the inter-cluster distance is greater than intra-cluster distance. The reported silhouette score was calculated as the average of all positive-valued Silhouette Coefficient trials^29^. A distribution was generated by calculating silhouette scores across many iterations (*n* = 50) of projection to 2D t-SNE space. A chance distribution was generated by repeating the silhouette score analysis for latent dynamics with randomly permuted articulator labels. Clustering with additional metrics is shown in Fig. S4.

### Alignment of latent dynamics

Single unaligned latent dynamic trajectories were visualized for S1 and S2 by plotting the first two PC values at each timepoint during the speech utterance window, averaged across all trials (Fig. 2c). We refer to these patient-specific latent dynamics as “unaligned”, as there has been no explicit functional alignment of these potentially different patient latent spaces to improve similarity between them. Alignment of these latent dynamics was performed using canonical correlation analysis (CCA) to find linear combinations of the latent dynamics PCs that resulted in maximal correlation between the latent dynamics of the two patients. CCA alignment is performed in a similar fashion as in^25^, which is described below and generally follows MATLAB’s *canoncorr* function. Alignment of latent dynamics was performed in Python 3.10 with numpy.

Starting with the latent dynamics of patients S1 (*s_1_, t, k_1_)* and S2 (*s_2_, t, k_2_)*, we require correspondence between trials *s_1_* and *s_2_*across these patients. As patients may be missing different trials due to experimental noise or outlier removal, we enforced uniformity in trials by condition averaging the trials into categories representing the unique phoneme sequences (*s_ca_,* at most 52 unique sequences) presented during the task and sub-selecting only the phoneme sequences present in both patients (i.e. if S1 has no trials for the sequences */abe/*, trials for */abe/* in S2 will not be considered for alignment). Condition-averaged trials were folded together with the time dimension to create latent dynamics matrices *L_1_, L_2_* of dimensions (*s_ca_*t, k_1_*) and (*s_ca_*t, k_2_*), respectively. We chose to condition average by phoneme sequence to enforce that the same phoneme sequences should have similar latent dynamics trajectories. We enforce this constraint by organizing *L_1_* and *L_2_* such that the same row/sample in each matrix corresponds to the same condition and timepoint. That is, we perform a single CCA fit between each pair of patients, where our condition constraint is enforced by matrix construction of *L_1_* and *L_2_*. Note that while label information is used to condition-averaged trials to create correspondence across patients, label information is not used in the following learning of CCA transformations to map between patient latent spaces.

Let *d* be the minimum rank of *L_1_* and *L_2_.* CCA alignment first centers *L_1_* and *L_2_* by mean subtraction. Following, *QR* decompositions are calculated for both *L*_1_ and *L_2_* as

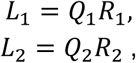

where *Q* is an orthonormal basis for the corresponding latent dynamics. Correlations between the orthonormal bases are represented by the inner product matrix *Q*_1_*^T^Q*_2_. Singular value decomposition on this matrix yields

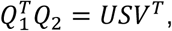

where *U* and *V* are the left and right singular vectors, respectively, which define the transformations of *Q_1_* and *Q_2_* to a shared, cross-patient space with maximal pairwise correlation. The *U* and *V* matrices (truncated to sizes (*k_1_, d*) and (*k_2_, d*), respectively) can be better understood in the context

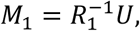

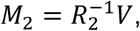

where *M_1_* and *M_2_* are projection matrices mapping *L_1_* and *L_2_* to the shared, cross-patient space, respectively. Transformation is performed via the matrix multiplication

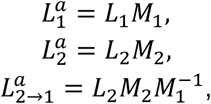

where *L*_1_*^a^ and L*_2_*^a^* are the aligned latent dynamics of S1 and S2, respectively, where the canonical correlations between these are the diagonal elements of *S* above. *L*_1_*^a^ and L*_2_*^a^* are aligned in a cross-patient space that requires transformations of both input patients. In this work, we instead choose to align patients to the space of a desired target patient to enable alignment from any number of target patients to a single common space. The alignment of S2’s latent dynamics to S1’s space, denoted *L*_2→1_*^a^*, is calculated by first aligning S2 to the common cross-patient space and applying the inverse of S1’s projection matrix (*M_1_*) to map to S1’s space. As such, multiple patients can be aligned to S1’s space by first calculating pairwise CCA alignment and projecting all patients to S1’s target space.

The clustering analysis with silhouette score quantification mentioned above was repeated on a dataset composed of both S1’s and S2’s data (Fig. 2d). Without alignment, latent dynamics were separately extracted from S1 and S2, concatenated (truncating the number of PCs in the patient with higher-dimensional latent dynamics to ensure consistency in dimensions for concatenation), and reduced to a 2D space with t-SNE. While we refer to these latent dynamics as “unaligned”, referencing a lack of explicit functional alignment between patients, the latent dynamics are effectively “variance-aligned”, in that there is assumed correspondence between PCs that explain the same relative levels of variance with each patient-specific latent space (e.g. PC 1 in S1 is assumed to correspond with PC 1 in S2). With alignment, S2 latent dynamics were aligned to S1’s space, aligned latent dynamics were concatenated, and t-SNE decomposition to a 2D space was applied. In both cases, silhouette score quantification was identical to the patient-specific case.

Representational similarity analysis (RSA)^55^ between unaligned and aligned patients was performed to quantify shared representations of speech production in aligned latent dynamics across all patients. For a given patient, we first construct a representational dissimilarity matrix (RDM). The latent dynamics from that patient (shape=(*s, t, k*)) are grouped by condition (articulator sequences in this analysis, e.g. “labial-low-dorsal” representing a sequence of three articulators) and averaged within these conditions to create a 2D data matrix of shape (*s_ca_, t*k*), where latent components and time have been folded together to a single dimension representing latent dynamics. For each pair of conditions, we calculate the Pearson correlation (*r*) between these flattened representative latent dynamics and represent their dissimilarity as 1 – *r.* The RDM is then constructed from these pairwise dissimilarities between different conditions within a patient (Fig. S6). Between a given source and target patient, each with RDMs constructed from latent dynamics, we calculate the representational similarity between the latent dynamics of these two patients as the correlation between their RDMs, indicating similarities in how articulatory speech information is represented in their respective latent dynamics. For each target patient, we quantify the average representational similarity between all source patients when RDM construction is performed using unaligned, patient-specific latent dynamics and latent dynamics aligned from the source patient to the target patient (Fig. 2f). We additionally calculated representational similarity when aligning surrogate data that is random with respect to speech (see *Generation of surrogate data)* and when labels were randomly permuted before RDM calculation (Fig. S7). This RSA analysis was performed in Python 3.10 using the *rsatoolbox* package (https://github.com/rsagroup/rsatoolbox).

### Single-electrode articulator tuning

Univariate tuning of electrodes to different articulator types was performed similar to Duraivel et al. (2023)^29^ (Fig. 3a). Briefly, for each patient and each electrode in that patient’s μECoG array, we trained a PCA-LDA (linear discriminant analysis) decoding model to predict articulator type from the timeseries HG data for the current electrode. This model first projected input data to a low-dimensional subspace explaining some percentage of the variance with PCA, then learned to classify articulator type from low-dimensional features. Evaluation and training were performed with 5-fold nested cross-validation. The outer folds, splitting training and testing data, were used to measure performance as the receiver-operator-characteristic-area-under-the-curve (ROC-AUC) for the binary classification of each articulator type. ROC-AUC scores were average across all 5-test folds within articulator type. The inner folds, splitting training and validation data, were used to optimize the PCA variance percentage hyperparameter. Grid search was performed on a one-dimensional vector of PCA variance percentages (10% to 100%, 5% step size) and the best performing percentage across all validation folds was used in the current test set fold. Maps for each articulator type (trial counts: low = 12, high=14, labial=14, dorsal=12). and patient were generated by repeating this process for all individual electrodes. Single-electrode articulator tuning was performed in Python 3.10 with the scikit-learn library.

### Cross-patient projection

To project single-electrode articulator maps from a source patient’s space to a target patient’s space (Fig. 3b), we first learned decompositions from HG signals to latent dynamics with PCA and pairwise alignment with CCA for each patient as described above (*Alignment of latent dynamics*), but using HG signals from all channels instead of only significant channels. We then applied these transformations to spatially decompose, optionally align, and spatially reconstruct articulator maps across patients. We note that alignment is learned from patient HG signals, not ROC-AUC scores, presenting no conflict of data leakage when projecting cross-patient articulator maps composed of these ROC-AUC values. The following equation summarizes this cross-patient projection process,

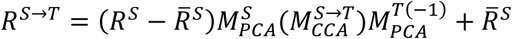

where 𝑅^S→T^ is the source patient’s articulator map projected to the target patient’s space, 𝑅^S^ is the source patient’s articulator map, 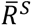 is the mean value of the source patient’s articulator map, 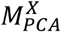 is the PCA decomposition for patient X (source or target), 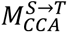 is the source-to-target CCA alignment matrix that can optionally be applied to perform cross-patient projection with alignment. When performing unaligned articulator map reconstructions, set 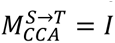. We also provide the following text description of this process for additional context. Decomposition was performed by passing flattened articulator maps through the source patient’s PCA. For compatibility between this PCA, which expects data centered around zero, and ROC-AUC values in the range [0.5, 1], we subtracted the mean ROC-AUC value across all channels to de-bias the ROC-AUC values. This enabled articulator maps to be scaled to proper feature ranges for transformation by the previously learned PCA. When aligned reconstructions were performed, alignment transformation matrices were applied to align the latent source patient articulator maps to the target patient’s latent space. With alignment, transformation to a different sized latent space is handled automatically. Without alignment, we enforced that each PCA used the same number of components (*n* = 30) instead of the number of components explaining 90% of the original variance for dimensionality consistency. Finally, the PCA transform from the target patient was inversely applied to map (optionally aligned) latent articulator maps up to the target patient’s electrode space. The mean ROC-AUC across all source patient channels (subtracted prior to transformation via source patient PCA) was then added back in to return ROC-AUC values to the proper domain. Fig. 3b (right) visualizes the reconstructions of S1 articulator maps by S3. Fig. S6 shows the reconstructed S1 articulator maps averaged across all source patients. We note that this reconstruction does not consider any neuroanatomical information in the projection. It is purely a reconstruction in electrode-level space, where we are interested in similarities between the reconstructed and ground-truth maps on an electrode-to-electrode level.

To compute the similarity between ground-truth and reconstructed articulator maps (Fig. 3c), the Pearson correlation was computed between ground-truth and reconstructed articulator maps. This is analogous to computing the cosine similarity between mean-subtracted ground truth and reconstructed articulator maps. Fig. 3c (left) quantifies the similarity for reconstructions of S1 articulator maps by only S3. In Fig. 3c (right), for each target patient, articulator maps from all source patients were considered by separately reconstructing these maps and averaging the maps across source patients as a single reconstructed map per articulator type prior to calculating similarity. We additionally calculated the correlation between reconstructed and ground-truth articulator maps when projecting cross-patient surrogate data (see *Generation of surrogate data*) that is random with respect to articulatory information (Fig. S9a).

We quantified the articulator decoding performance of models tested on aligned and unaligned reconstructed data (Fig. 3d). All decoding models used were PCA-SVM (support vector machine) models. This combined a PCA layer for feature size reduction (as in PCA-LDA models described above) with a bagged SVM classifier (scikit-learn, C=1, gamma=(n_features*feature_variance)^-1^, balanced class weight) using a radial basis function kernel. No additional feature scaling beyond PCA reduction was performed prior to SVM input. SVM PCA explained variance was set at a fixed 80% value. Articulator decoding performance (4 classes: low, high, labial, dorsal; chance = 0.25) was quantified as the balanced accuracy across predictions from all test folds with 20-fold cross-validation. The articulator identity for the first phoneme of the repeated three-phoneme sequence was predicted by the models. First, we trained baseline patient-specific decoding models for each patient, where both training and testing data were composed of data from a single patient. We then projected HG data from each source patient’s (S2-S8) to S1’s electrode space by inversely applying S1’s PCA transformation to the source patient’s latent dynamics, either unaligned or aligned to S1’s space. Cross-patient decoding was performed by applying the decoding model trained on S1’s data to each source patient’s reconstructed HG data as test data. We additionally performed cross-patient articulator decoding when projecting cross-patient surrogate data (see *Generation of surrogate data*) that is random with respect to articulatory information (Fig. S9b). Similar to above, cross-patient decoding was performed by applying the decoding model trained on S1’s real data, to each source patient’s surrogate data reconstructed in S1’s space as test data.

### Cross-patient phoneme decoding

We trained PCA-SVM decoding models (as described above in *Cross-patient projection*) to predict phoneme (9 classes: */a/, /e/, /i/, /u/, /b/, /p/, /v/, /g/, /k/*; chance = 0.11) identity in a target patient from combined cross-patient data across all patients in our dataset (Fig. 4a). Parameters and evaluation methods are identical to those defined above, but with phonemes as the labels instead of articulators. Phonemes were predicted at all three positions of the phoneme sequence. Phoneme decoding accuracy was reported as the balanced accuracy across predictions from all 20 test folds across 50 iterations to generate a distribution across different training and testing splits. Cross-validation folds were generated across the trial dimension of our data. For results aggregated across all patients (Fig. 4b), the average balanced accuracy across all 50 iterations was reported as a single value per patient. For all decoding contexts (patient-specific, unaligned cross-patient, aligned cross-patient), training and test splits were only performed on the target patient. That is, in cross-patient contexts, cross-patient data was only aligned to the target patient’s training data (when alignment was performed) and added to the training set, such that model accuracy was quantified on held-out data from a single target patient, as would be the clinical use-case for cross-patient speech BCIs. This alignment to only the training set of the target patient additionally prevented data leakage between the training and testing data. Only HG data from significant channels were used for cross-patient phoneme decoding. Cross-patient phoneme decoding was performed in Python 3.10, primarily with the scikit-learn library.

We started by calculating the phoneme decoding accuracies of patient-specific models for each target patient as a baseline. Models were trained on unaligned cross-patient data by first separately decomposing all patient HG data to latent dynamics explaining 90% of the variance of the input HG data with PCA. For the target patient, PCA decomposition was learned on the training set and applied to the test set to avoid information leakage. Unaligned latent dynamics across all patients were concatenated as additional observations/trials into a single dataset (truncated to the minimum number of PCs across all patients for consistency of latent dimensionality). A PCA-SVM decoding model was then trained to predict phoneme identity from the expanded cross-patient training dataset. As mentioned above, phoneme decoding performance of the PCA-SVM was quantified on the test set containing only data from the target patient. Models were trained on aligned cross-patient data by repeating the same process for unaligned cross-patient data and applying CCA alignment after decomposing all patient data to latent dynamics. CCA alignment was calculated between each source patient and the target patient to align all source patient data to the same target patient space (this was additionally beneficial as no additional transformation of the test data in the target patient’s space was required). As with PCA, CCA alignment was only learned between the training set of the target patient and cross-patient sources to prevent information leakage between training and testing sets. Fig. 4b shows cross-patient phoneme decoding results S1, S2, S3. The results across all patients are shown in Fig. S7.

As a control condition for cross-patient decoding, we aligned surrogate data that is random with respect to phoneme information (see *Generation of surrogate data*) to each target patient instead of true data from other patients (Fig. S11). Surrogate data were generated for each of the eight patients, where seven surrogate patients were aligned to true target patient data for each target patient. Alignment and subsequent decoding were done the same as with all true data.

For subsampling of target patient data (Fig. 4c, Fig. S9 for all patients separately), patient-specific and aligned cross-patient phoneme decoding were performed as described above while a percentage of the target patient’s training trials were subsampled prior to alignment (for cross-patient models) and decoding. Subsampling was performed by randomly and uniformly sampling trials from the target patient’s training dataset until the desired number of trials were obtained. Reported phoneme decoding accuracy is the average balanced decoding accuracy for each of the eight target patients.

For subsampling of cross-patient data (Fig. 4d, Fig. S10 for all patients separately), aligned cross-patient models were trained as described above while varying the amount of cross-patient data aligned to the target patient. We equally subsampled *n* trials from each patient, where *n* ranged from 5 trials to the maximum number of trials from the current patient with a step size of 25 (i.e. 5, 30, 55, …). Cross-patient trials were sampled equally from each patient to avoid effects of patient identity during sampling. Each point reports the phoneme decoding accuracy across 50 iterations of sampling. Fig. 4d visualizes these results for target patient S5. The average patient-specific decoding accuracy for S5 is overlaid for comparison. To show diminishing gains in decoding accuracy as the amount of cross-patient data increased, we performed a linear fit to S5’s log-transformed decoding accuracy vs. cross-patient trials curve using scipy (Fig. S11).

To investigate inter-patient variability in aligned cross-patient decoding, we quantified each target patient’s percent change in decoding accuracy from patient-specific performance to aligned cross-patient decoding with all source patients added, which denote as cross-patient improvement. We plotted this cross-patient improvement against the decoding accuracy of the target patient and found a significant relationship using a linear fit between the two variables (Fig. 4e). As mentioned in the main text, we chose to exclude S3 from this analysis and following inter-patient variability due to its small dataset size, as it would be unclear cross-patient data would disproportionately increase accuracy in this dataset with its limited number of trials. We additionally investigated inter-patient variability using a leave-one-out cross-patient analysis. Here, for a given target patient, we trained an aligned cross-patient model with all available source patients except one, which was parametrically varied as the left-out patient. Then we quantified the benefit of that left-out patient’s data to the target patient’s data as the percent change in accuracy between aligned cross-patient training with all source patients and aligned cross-patient training without the left-out patient, which we call the left-out accuracy change (Fig. S19a). Similar to Figure 4e, we regressed the left-out accuracy change against effects of 1) target patient-specific accuracy (Fig. S19b)., 2) left-out patient-specific accuracy (Fig. S19c)., and 3) distance between electrode arrays in neuroanatomical space (Fig. S19d). All relationships against left-out accuracy change were assessed with linear fits between the left-out accuracy change and the desired variable. P-values of each linear fit were FDR corrected over all fits performed in the leave-one-out analysis,

### Simulated real-time data generation

We simulated the real-time collection of neural features from our offline speech data to evaluate the effectiveness of our proposed alignment method in contexts more applicable to speech BCI deployment. To accomplish this, we adapted the methods of Willett and colleagues^7^ to our high-gamma processing and phoneme decoding pipeline. We began with 2ksps neural data split into trials by alignment to the presentation of auditory stimuli during the speech task (as real-time methods would not have knowledge of patient response times to use response-aligned data). We split each trial into 20 ms bins (40 timepoints per bin at 2 ksps), applied a common average reference within bins, and performed a modified HG filtering method within these short time bins. HG filtering methods for offline, trial-based data typically employ a Hilbert transform to extract the envelope of the logarithmically-spaced and sized frequency bands in the HG range (see *High-gamma extraction)*, which can be difficult to accurately implement in low-latency contexts with short processing bins. Because of this, we chose to calculate the root-mean-square (RMS) HG power in each of these 20 ms bins instead. We again bandpass the signal into 8 logarithmically-spaced and sized frequency bands as with our offline data, then calculate RMS power in each band as simply the square root of the mean of the squared signal. Bandpass filtering was performed causally using a 4^th^ order Butterworth filter designed to match frequency band characteristics of the offline filtering frequency bands^50^. We finally average across these bands to calculate a single HG power value for each channel. This is repeated for every 20 ms bin within each trial, effectively downsampling the output HG power to a sampling rate of 50 sps. Here, trials were initially defined 0 s to 3.5 seconds relative to stimulus onset. However, we trimmed the initial 500 ms to remove filter artifacts, resulting in trial definition as 0.5 s to 3.5 s relative to stimulus onset.

To mimic clinical speech BCI deployment contexts with designated training and evaluation periods, we split our three-block speech task into training blocks (first two blocks) and a testing block (last block). Due to the single block of data collection for patient S3, we chose to exclude them as a target patient in our simulated real-time decoding analyses (though we still used their data to increase training set size in other target patients). Following simulated real-time processes of the data from the training period, we performed channel and trial outlier removal identically to our offline methods (see *Neural data preprocessing*). Identified bad channels from the training period were saved and excluded from the data in the test period by imputing their values with the mean across all good channels. Real-time processed HG data aligned to the stimulus were normalized (mean-subtraction) by a pre-stimulus baseline (-500 ms to 0 ms relative to stimulus onset as with offline data). Due to the short time period of experimental data collection, we chose to save this baseline HG data from the training period of normalization of the test data as well instead of using rolling normalization methods present in studies with longer collection and evaluation periods.

### Connectionist temporal classification recurrent neural network (CTC-RNN) decoding

Following simulated real-time processing of our training period and testing period data, we trained stacked recurrent neural network (RNN) models with connectionist temporal classification loss (CTC) to continuously predict phoneme identity throughout the trials. We used CTC loss and decoded stimulus-presented phonemes (instead of patient response phonemes), as in previous speech BCI work^7,8^, to avoid use of information on patient-response timing which would not be available in real-time speech BCI settings. Specifically, we set the label for each speech trial as the sequence of phonemes presented in the auditory stimulus. Then from 0.5 s to 3.5 s post-stimulus onset, we decoded phoneme identity using a causal sliding window of HG power features (280 ms window size, 80 ms stride, i.e. 14 sample window size, 4 sample stride) as performed by Willett and colleagues^7^. Training with CTC loss enables the comparison of different length sequences in “decoding window” space (predicted phoneme every 80 ms) to “stimulus sequence” space (three-length sequence of phonemes) by collapsing repeatedly predicted phonemes and ignoring periods of “blank” predictions with no associated phoneme. In patient-specific decoding cases, we trained models to directly translate real-time processed HG data across all channels to phoneme sequences via our stacked RNN model with CTC loss. In unaligned cross-patient decoding cases, high-channel count data was first reduced to patient-specific latent spaces via PCA, pooled together into a single training set across patients, the trained to predict phoneme sequences with the stacked RNN model. To enable data pooling across different latent sizes, latent spaces were truncated to the minimum latent size across all patients. Latent size was determined by the number of components explaining 90% of the variance in the input data. However, for patient S5, noise in a single channel that was not removed by our initial channel removal resulted in a single component explaining greater than 90% of the data’s variance and poor subsequent alignment. To remedy this, we fixed component selection to 30 components to extract a broader range of components that would be better suited for subsequent alignment in this patient. As latent space reduction was performed on data from the training period, this manual removal is still valid in the context of clinical speech BCI deployment, as a similar analysis and correction of data could be performed between training and testing periods. In aligned cross-patient decoding cases, decoding performed as in the unaligned case, but with alignment of all source patient latent dynamics to the target patient space prior to model training. In both unaligned and aligned cases, learned dimensionality reduction and alignment linear transformations were saved for application to validation or test set data.

CTC-RNN models were trained to optimize phoneme error rate (PER) between true phoneme sequences and predicted phoneme sequences. PER was defined as the number of insertions, substitutions, or deletions necessary to convert the predicted sequence to the true sequence, divided by the true sequence length and converted to a percentage. In all decoding contexts (patient-specific, unaligned, aligned), we trained the model over 300 epochs. We applied the following data augmentations to our training data:

– **Temporal warping**: Each trial was temporally warped by randomly stretching or compressing the time axis using linear interpolation. The signal was first resampled to a randomly scaled duration (factor range: 0.8–1.2) and then resampled back to the original length, introducing local temporal distortions while preserving overall sequence duration.
– **Temporal masking**: A contiguous segment of each trial (10% of the total duration) was randomly selected and set to zero, simulating transient signal dropout and encouraging robustness to missing temporal information.
– **Time shifting**: Each trial was circularly shifted along the time axis by a random offset (±20 samples), preserving signal structure while introducing temporal misalignment to improve invariance to onset timing.
– **Additive noise**: Additive Gaussian noise (σ = 0.01) was applied to the signal to simulate measurement noise and improve robustness to low-amplitude perturbations.
– **Amplitude scaling**: Each trial was multiplied by a random scalar factor (range: 0.9–1.1), introducing variability in overall signal amplitude to promote invariance to differences in amplitude.

We split a validation set from our training data to monitor PER during training and loaded the model weights from the model with lower PER during training for use in evaluation on the test set (Fig. 6b). We performed hyperparameter optimization for each patient and decoding context separately to optimize CTC-RNN performance. Hyperparameter optimization was implemented with random search and 10-fold cross-validation over training trials (see Tables S6-8 for patient hyperparameters in patient-specific, unaligned, and aligned cases, respectively). As the sequence length of predicted phoneme sequence is variable, a theoretical chance value of PER based on phoneme composition is less defined for this analysis. As such, we calculated an empirical chance distribution for each patient by performing patient-specific CTC-RNN training with permuted labels and evaluation on unmodified test data. As with our offline decoding analysis, we additionally performed CTC-RNN decoding evaluations while subsampling the amount of target patient data (Fig. 5c) and cross-patient data (Fig. 5d). We projected PER with larger cross-patient data amounts by performing a linear fit between aligned decoding PER and log-transformed amounts of cross-patient data using scipy’s *linregress* function. All CTC-decoding analyses were performed in Python 3.10 via Pytorch-lightning v2.3.3 and Pytorch v2.4.0.

### Pitch subsampling

Lower electrode densities were simulated by using Poisson disk sampling^56^ to sub-select electrodes from the μECoG array constrained by a minimum distance between electrodes. Given a desired pitch as this constraining distance, we calculated the number of electrodes that needed to be sampled to achieve this pitch as:

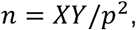

where *n* is the number of electrodes, *X* and *Y* are the length and width of the μECoG array, and *p* is the desired pitch. We rounded *n* to the nearest integer in implementation. We only considered subsampled electrodes with significant HG activity during the patient response for subsequent cross-patient analyses. In these cross-patient analyses, all patients were subsampled to the desired pitch, not just the target patient.

Cross-patient decoding on pitch-subsampled data (Fig. 5a, top; Fig. S13 for all patients separately) was performed as described previously (*Cross-patient phoneme decoding*) with PCA-SVM models, but with pitch subsampling of the HG data prior to latent dynamics extraction, alignment, and decoding. At each of the 50 training iterations, a new subsampling of electrodes at the current pitch was selected to mitigate effects of electrode identity.

Representational similarity between cross-patient latent dynamics from subsampled arrays (Fig. 5a, bottom) was calculated similarly to previous analyses (*Alignment of latent dynamics*) except at each pitch, the representational similarity was averaged across 50 iterations to mitigate effects of electrode identity during sampling.

### Grid subsampling

Smaller electrode array coverages were simulated by sampling rectangular sub-grids of size *n* x *2n* (*n ɛ [2,8]*) from the total electrode array. While maximum electrode array sizes exceeded 8x16 for patients with arrays of 256 electrodes, we only considered sub-grids up to 8x16 for consistency across all patients. For a given sub-grid size, we consider all possible starting locations that fit within the total electrode array. That is, for an 8x16 electrode array with a sub-grid size of 6x12, there would be a total of 15 possible sub-grids to select from. Only electrodes with significant HG activity during the patient response were selected from each sub-grid.

Cross-patient decoding from sub-grids (Fig. 5b, top; Fig. S14 for all patients separately) was again performed as previously described with PCA-SVM models and followed a similar approach to the cross-patient correlation analysis for sub-grid selection. For a given sub-grid size, training was repeated for all possible target patient sub-grids and a random sub-grid from each source patient was selected at each iteration. At each sub-grid size, phoneme decoding accuracy was averaged across all target patient sub-grids. Representational similarity between cross-patient latent dynamics from sub-grids (Fig. 5b, bottom) were extracted similarly to previous analyses. For each target patient and sub-grid size, we extracted latent dynamics from HG in the sub-selected electrodes and aligned them across patients. We iterated over all possible sub-grids of the given size for the target patient. For each possible sub-grid, we selected a random sub-grid (uniformly) from the possibilities for each source patient. Following, representational similarity across aligned latent dynamics was calculated as previously described. For each sub-grid size, the reported correlation was averaged across all target patient sub-grids and across all source patients, yielding a single averaged representational similarity value for each target patient.

### Spatial-averaging

Larger contact sizes were simulated by averaging data together from neighboring contacts with varying sizes. Contact sizes ranged from 1x1 (unaveraged micro-contact used in previous analyses in this work) to 8x8 squares. To simulate neural recording with these larger contacts, raw signals were averaged together on these contacts and HG was extracted from the resultant spatially-averaged neural signal. Only spatially-averaged contacts with significant HG activity during the patient response (relative to a baseline period 500 ms prior to stimulus presentation) were considered for subsequent analyses (see *Statistical Analysis*). Note that this spatial-averaging method does not account for changes in electrode impedance for different sized contacts. For contact sizes that did not span the total electrode array (e.g. Two 8x8 contacts span an 8x16 electrode array, but 7x7 contacts do not), electrodes were centered within the electrode array.

Cross-patient decoding on spatially-averaged data (Fig. 5c, top; Fig. S15 for all patients separately) was performed as described previously (*Cross-patient phoneme decoding*) with PCA-SVM models, but with the spatially-averaged data used for latent dynamics extraction, alignment, and decoding. Cross-patient representational similarity in latent dynamics for spatially averaged data (Fig. 5c, bottom) was again calculated similar to previous analyses, but with spatial-averaging of data prior to latent dynamics extraction and alignment. As the spatial-averaging results in a singular, deterministic sub-selection (unlike pitch and grid subsampling), the representational similarity in latent dynamics was not repeated over multiple iterations. Instead, a single representational similarity value (Pearson *r* between RDMs) was calculated of each target patient and source patient and averaged across all source patients for each target patient (as in *Alignment of latent dynamics*). For each contact size, representational similarity was reported across a distribution of all target patients.

### Statistical analysis

All statistical analyses were performed in MATLAB R2021a and Python 3.10 (scipy and statsmodels). All permutation tests mentioned below use the difference in means as the test statistic. All false discovery rate (FDR) control mentioned below implements the Benjamini-Hochberg method^57^.

## Data Availability

The data from µECoG recordings that support the decoding analysis are available via the Data Archive for The Brain Initiative (DABI; accession code: <TO be updated after acceptance>) under restricted access (restrictions will be exempted after appropriate modifications of the IRB protocol: Pro0072892). The access can be obtained upon written requests from the corresponding authors. Source data are provided with this paper.

## Code Availability

Code for cross-patient alignment analyses and notebooks recreating the figures in this work can be found at https://github.com/coganlab/cross_patient_speech_decoding.

## Acknowledgements

J.V., G.B.C., S.P.L., A.H.F., D.G.S, S.D., Z.S. were supported by NIH R01DC019498 and DoD W81XWH-21-0538. S.R. was supported by an NRCDP NIH K12 award. J.V., G.B.C., C.W., S.D. were supported by NIH CTSA UL1TR002553. K.B. was supported by the National Science Foundation (Graduate Research Fellowship Program DGE-1644868). C.S. was supported by the National Science Foundation (Graduate Research Fellowship Program DGE-2139754) J.V. and G.B.C. were supported by NIH UG3NS120172, G.B.C, J.V., S.P.L, A.H.F, D.G.S. were supported by NIH R01NS129703, and G.B.C., J.V., S.P.L., S.D. were supported by an internal Duke Institute for Brain Sciences Incubator Award. We thank Nicole Liddle, Palee Abeyta, and Anna Thirakul for help with consenting participants, Seth Foster for help with the experimental design, Steve Harward for help with surgical preparation, and Daniel Sexton for help with surgical preparation and neuroanatomical localizations of electrode arrays. With tremendous gratitude, we thank the patients who volunteered their time for our study.

## Author Contributions

Z.S. performed all analyses, performed statistical testing, and generated figures. S.D. performed significant preprocessing of the neural data and assisted in analysis design. G.B.C. and S.R. designed the experimental task. S.R., S.P.L., A.H.F, D.G.S performed intraoperative electrode implantation and neural data acquisition. C.W., K.B., C.S., and J.V. were involved in µECoG electrode design and setting up the intraoperative data acquisition process. Z.S., S.D, K.B., C.S., and J.V., and G.B.C performed intraoperative neural data collection. Z.S., J.V., and G.B.C wrote the manuscript with feedback from other co-authors. G.B.C. was involved in clinical supervision. J.V., and G.B.C. designed and supervised the human intraoperative study.

