## Supplementary Figures and Tables for "Shared latent representations of speech production for cross-patient speech decoding"

### Supplementary Tables

**Table S1. Clinical summary of patients**

| Patient | Age | Sex | Diagnosis | MDS-UPDRS Part III Speech Score* | Number of channels |
|---|---|---|---|---|---|
| S1 | 61 | M | Parkinson’s | 2 (Mild) | 128 |
| S2 | 64 | M | Parkinson’s | 0 (Normal) | 128 |
| S3 | 26 | F | Tumor resection | N/A | 256 |
| S4 | 63 | M | Parkinson’s | 2 (Mild) | 128 |
| S5 | 61 | M | Parkinson’s | 1 (Slight) | 128 |
| S6 | 71 | M | Tumor resection | N/A | 256 |
| S7 | 53 | M | Tumor resection | N/A | 256 |
| S8 | 60 | M | Tumor resection | N/A | 256 |

**Mean age:** 57.4

***Movement Disorder Society-Unified Parkinson's Disease Rating Scale (MDS-UPDRS) Part III** is a clinical standard for evaluating motor impairment of patients with Parkinson’s disease. Scores listed here show evaluation of speech-related motor impairment. Scores range from 0-4 with 0=Normal, 1=Slight, 2=Mild, 3=Moderate, 4=Severe).

**Table S2. Task stimuli**

| CVC | VCV |
|---|---|
| */bab/* | */abe/* |
| */bek/* | */abi/* |
| */bak/* | */eba/* |
| */bup/* | */ebi/* |
| */gab/* | */ebu/* |
| */geb/* | */ega/* |
| */gev/* | */eka/* |
| */gak/* | */epi/* |
| */gav/* | */aka/* |
| */gig/* | */aku/* |
| */gip/* | */ava/* |
| */gub/* | */ave/* |
| */kab/* | */ibu/* |
| */keg/* | */ika/* |
| */kub/* | */ike/* |
| */kug/* | */ipu/* |
| */pek/* | */iva/* |
| */pep/* | */ivu/* |
| */pev/* | */uba/* |
| */puk/* | */uga/* |
| */pup/* | */uge/* |
| */vek/* | */uke/* |
| */veg/* | */upi/* |
| */vip/* | */upu/* |
| */vug/* | */uve/* |
| */vuk/* | */uvi/* |

**Table S3. Patient data summary**

| Patient | No. of trials with patient response | Approx. utterance duration (min.) | Approx. experiment duration (min.) | No. of channels with significant HG activity (no. and % of total) |
|---|---|---|---|---|
| S1 | 144 | 1.08 | 9.00 | 111/128 (86.7%) |
| S2 | 148 | 1.11 | 9.25 | 111/128 (86.7%) |
| S3 | 46 | 0.35 | 2.88 | 149/256 (58.2%) |
| S4 | 151 | 1.13 | 9.44 | 74/128 (57.8%) |
| S5 | 151 | 1.13 | 9.44 | 63/128 (49.2%) |
| S6 | 137 | 1.03 | 8.56 | 144/256 (56.3%) |
| S7 | 141 | 1.06 | 8.81 | 171/256 (66.8%) |
| S8 | 178 | 1.34 | 11.13 | 201/256 (78.5%) |

**Total utterance duration across patients:** ~8.23 minutes
**Total experiment duration across patients:** ~68.51 minutes

**Table S4. Latent space dimensionality**

| Patient | Significant channels | Latent components (*k*, 90% variance explained) |
|---|---|---|
| S1 | 111 | 25 |
| S2 | 111 | 26 |
| S3 | 149 | 43 |
| S4 | 74 | 22 |
| S5 | 63 | 20 |
| S6 | 144 | 65 |
| S7 | 171 | 54 |
| S8 | 201 | 67 |

**Table S5. Pairwise shared latent space dimensionality**

| Target patient | Source patient | Shared latent components (*d*) |
|---|---|---|
| S1 | S2 | 25 |
| S1 | S3 | 25 |
| S1 | S4 | 22 |
| S1 | S5 | 20 |
| S1 | S6 | 25 |
| S1 | S7 | 25 |
| S1 | S8 | 25 |
| S2 | S3 | 26 |
| S2 | S4 | 22 |
| S2 | S5 | 20 |
| S2 | S6 | 26 |
| S2 | S7 | 26 |
| S2 | S8 | 26 |
| S3 | S4 | 22 |
| S3 | S5 | 20 |
| S3 | S6 | 43 |
| S3 | S7 | 43 |
| S3 | S8 | 43 |
| S4 | S5 | 20 |
| S4 | S6 | 22 |
| S4 | S7 | 22 |
| S4 | S8 | 22 |
| S5 | S6 | 20 |
| S5 | S7 | 20 |
| S5 | S8 | 20 |
| S6 | S7 | 54 |
| S6 | S8 | 65 |
| S7 | S8 | 54 |

* *d* is the same for each pair of patients, regardless of which is the target and source, so redundant rows have been omitted.

**Table S6. CTC-RNN hyperparameters (patient-specific)**

| | Patient | | | | | | |
|---|---|---|---|---|---|---|---|
| | S1 | S2 | S4 | S5 | S6 | S7 | S8 |
| **Batch size*** | 128 | 256 | 256 | 128 | 256 | 128 | 256 |
| **Kernel size** | 280 ms (14 samples) | 280 ms | 280 ms | 280 ms | 280 ms | 280 ms | 280 ms |
| **Stride** | 80 ms (4 samples) | 80 ms | 80 ms | 80 ms | 80 ms | 80 ms | 80 ms |
| **RNN unit type** | GRU | GRU | GRU | GRU | GRU | GRU | GRU |
| **# RNN units (hidden size)*** | 128 | 256 | 256 | 128 | 256 | 128 | 256 |
| **# RNN layers*** | 5 | 4 | 5 | 4 | 5 | 5 | 5 |
| **RNN dropout*** | 0.4 | 0.3 | 0.2 | 0.3 | 0.4 | 0.3 | 0.3 |
| **Total epochs**** | 300 | 300 | 300 | 300 | 300 | 300 | 300 |
| **Learning rate*** | 1e-3 | 5e-4 | 5e-3 | 5e-3 | 5e-3 | 1e-3 | 5e-3 |
| **L2 Regularization*** | 1e-3 | 1e-4 | 1e-3 | 1e-3 | 1e-3 | 1e-5 | 1e-4 |
| **Optimization** | AdamW | AdamW | AdamW | AdamW | AdamW | AdamW | AdamW |
| **Gradient clipping** | 5.0 | 5.0 | 5.0 | 5.0 | 5.0 | 5.0 | 5.0 |

* Tuned hyperparameters
** While 300 total epochs were run for training, model weights from the checkpoint with the highest accuracy on the validation set were loaded for use on the test set.

**Table S7. CTC-RNN hyperparameters (unaligned)**

| | Patient | | | | | | |
|---|---|---|---|---|---|---|---|
| | S1 | S2 | S4 | S5 | S6 | S7 | S8 |
| **Batch size*** | 256 | 256 | 256 | 128 | 128 | 128 | 128 |
| **Kernel size** | 280 ms (14 samples) | 280 ms | 280 ms | 280 ms | 280 ms | 280 ms | 280 ms |
| **Stride** | 80 ms (4 samples) | 80 ms | 80 ms | 80 ms | 80 ms | 80 ms | 80 ms |
| **RNN unit type** | GRU | GRU | GRU | GRU | GRU | GRU | GRU |
| **# RNN units (hidden size)*** | 512 | 256 | 256 | 512 | 512 | 512 | 512 |
| **# RNN layers*** | 5 | 5 | 4 | 5 | 4 | 3 | 4 |
| **RNN dropout*** | 0.3 | 0.2 | 0.4 | 0.3 | 0.4 | 0.2 | 0.3 |
| **Total epochs**** | 300 | 300 | 300 | 300 | 300 | 300 | 300 |
| **Learning rate*** | 1e-3 | 1e-3 | 5e-3 | 5e-4 | 5e-3 | 5e-3 | 5e-3 |
| **L2 Regularization*** | 1e-4 | 1e-5 | 1e-3 | 1e-3 | 1e-4 | 1e-4 | 1e-3 |
| **Optimization** | AdamW | AdamW | AdamW | AdamW | AdamW | AdamW | AdamW |
| **Gradient clipping** | 5.0 | 5.0 | 5.0 | 5.0 | 5.0 | 5.0 | 5.0 |

* Tuned hyperparameters

** While 300 total epochs were run for training, model weights from the checkpoint with the highest accuracy on the validation set were loaded for use on the test set.

**Table S8. CTC-RNN hyperparameters (aligned)**

| | Patient | | | | | | |
|---|---|---|---|---|---|---|---|
| | S1 | S2 | S4 | S5 | S6 | S7 | S8 |
| **Batch size*** | 128 | 128 | 256 | 128 | 256 | 128 | 128 |
| **Kernel size** | 280 ms (14 samples) | 280 ms | 280 ms | 280 ms | 280 ms | 280 ms | 280 ms |
| **Stride** | 80 ms (4 samples) | 80 ms | 80 ms | 80 ms | 80 ms | 80 ms | 80 ms |
| **RNN unit type** | GRU | GRU | GRU | GRU | GRU | GRU | GRU |
| **# RNN units (hidden size)*** | 512 | 512 | 256 | 256 | 512 | 256 | 512 |
| **# RNN layers*** | 4 | 4 | 4 | 4 | 5 | 5 | 5 |
| **RNN dropout*** | 0.4 | 0.4 | 0.4 | 0.4 | 0.3 | 0.4 | 0.4 |
| **Total epochs**** | 300 | 300 | 300 | 300 | 300 | 300 | 300 |
| **Learning rate*** | 1e-3 | 1e-3 | 5e-3 | 1e-3 | 1e-3 | 5e-3 | 5e-4 |
| **L2 Regularization*** | 1e-3 | 1e-3 | 1e-3 | 1e-4 | 1e-4 | 1e-4 | 1e-4 |
| **Optimization** | AdamW | AdamW | AdamW | AdamW | AdamW | AdamW | AdamW |
| **Gradient clipping** | 5.0 | 5.0 | 5.0 | 5.0 | 5.0 | 5.0 | 5.0 |

* Tuned hyperparameters
** While 300 total epochs were run for training, model weights from the checkpoint with the highest accuracy on the validation set were loaded for use on the test set.

### Supplementary Figures

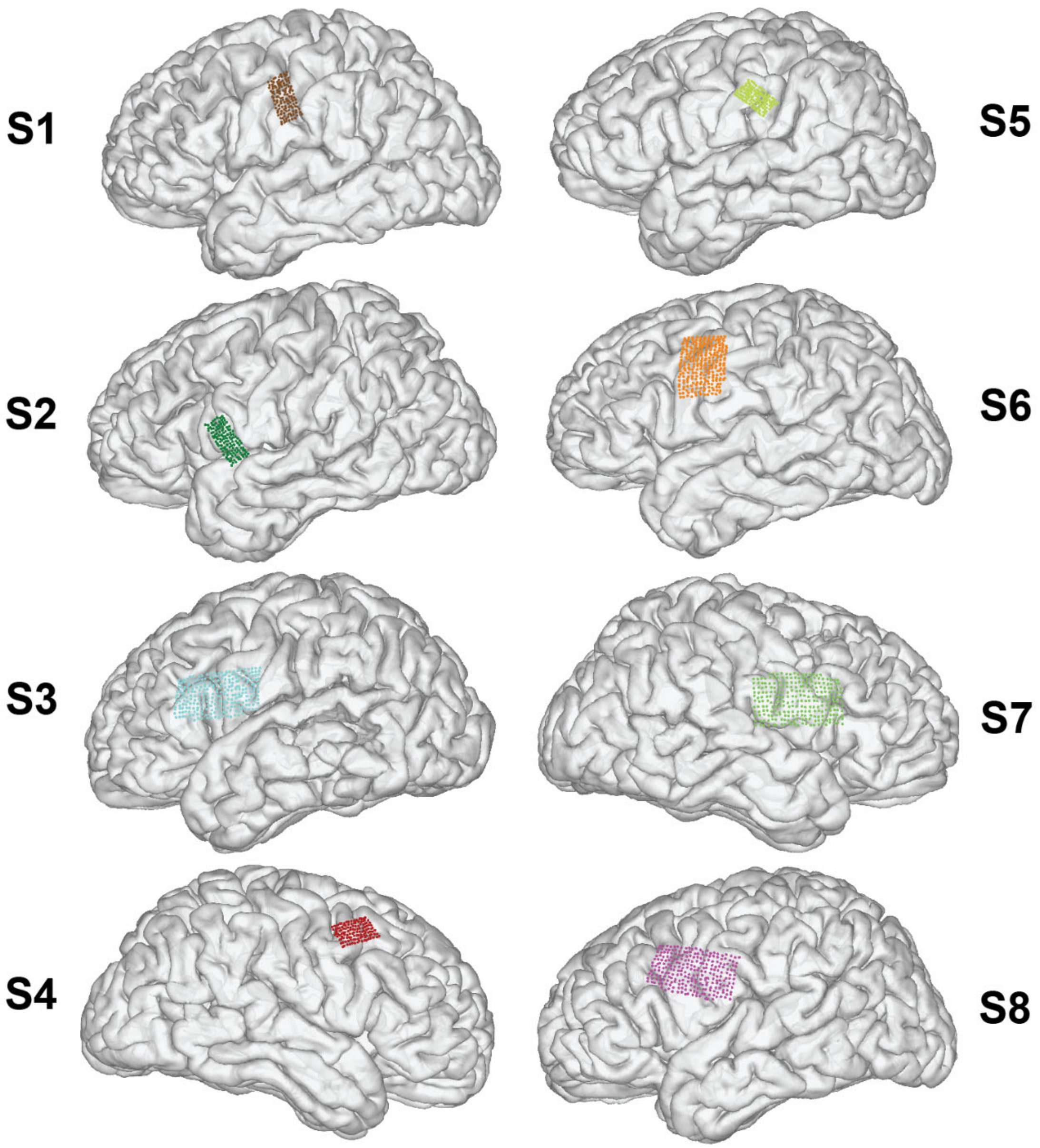


**Figure S1. Patient-specific electrode array locations.** Neuroanatomical locations of µECoG arrays as shown in Fig. 1, but on patient-specific brains instead of projected to a common brain.

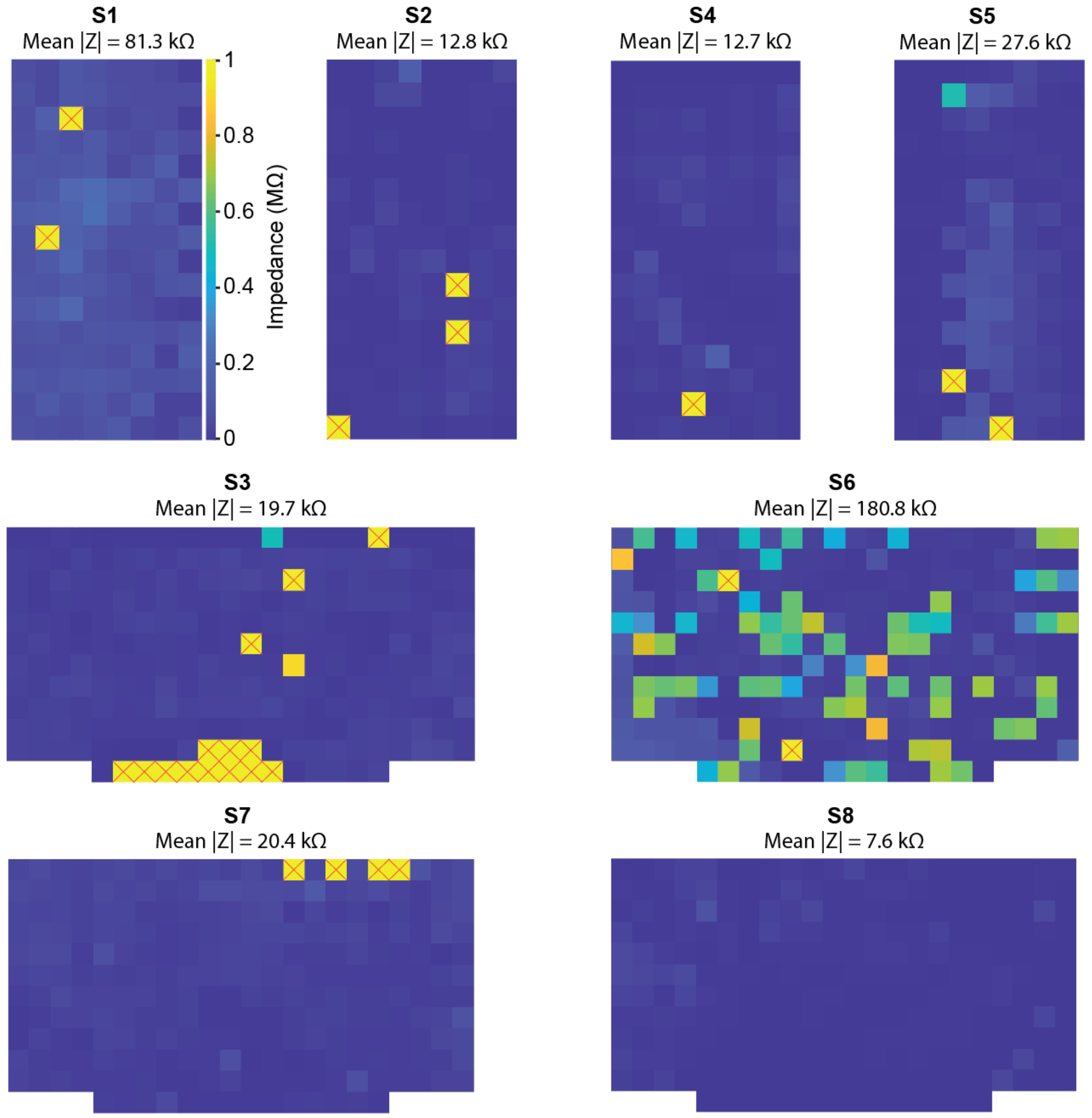


**Figure S2. Electrode impedance maps.** Impedance values (MOhm) from each patient's electrode array. Channels with a red X indicate channels with impedance values above 1 MOhm that are marked as bad channels in our preprocessing pipeline. All electrodes show low impedance values except in S6.

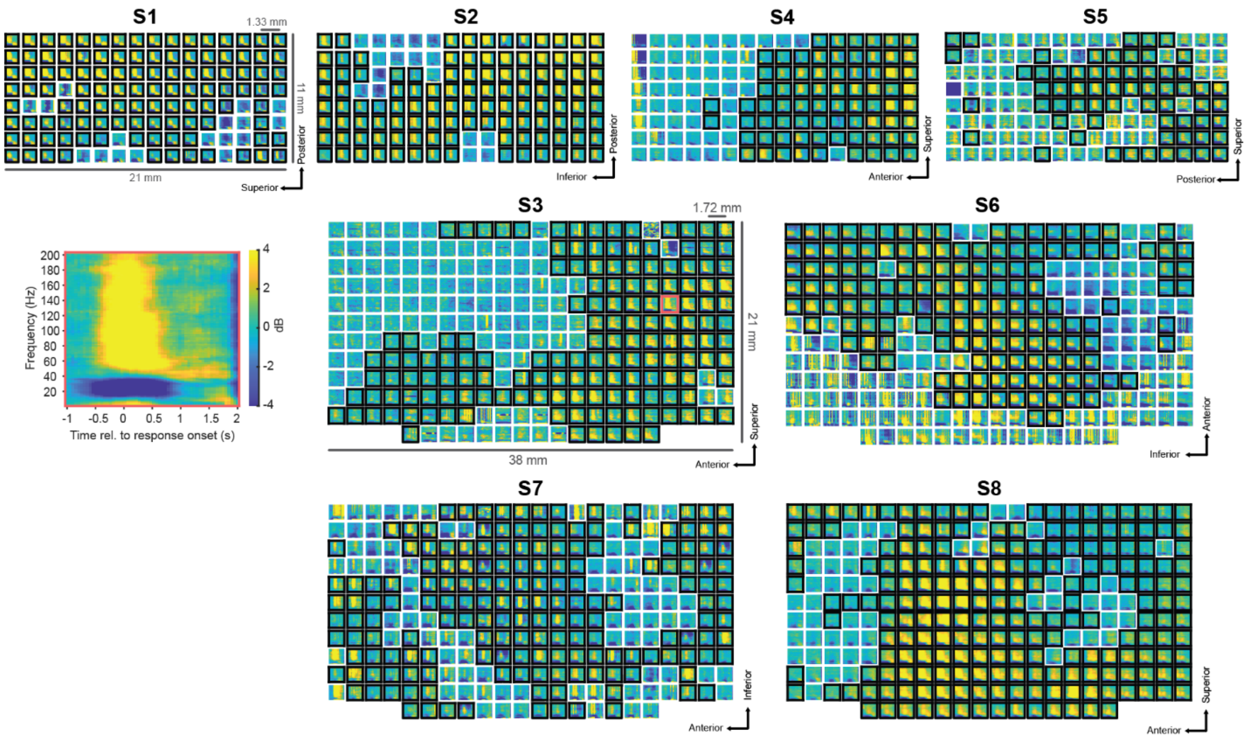


**Figure S3. Spectrotemporal speech activations.** Spectrotemporal activation patterns for all electrodes across the µECoG arrays of all patients (S1-S8), where each box represents a single electrode. Electrodes with black borders were found to have significant HG activity during the patient response, relative to a pre-stimulus baseline (see Methods). The spectrogram on the left shows a single channel from S3's array with relevant HG activity around speech production. µECoG array characteristics (size, pitch) are annotated on the S1 (128 channels) and S3 (256 channels) arrays. Axes on the bottom right of each array indicate the neuroanatomical orientation of the array (See Fig. S1 for full neuroanatomical locations).

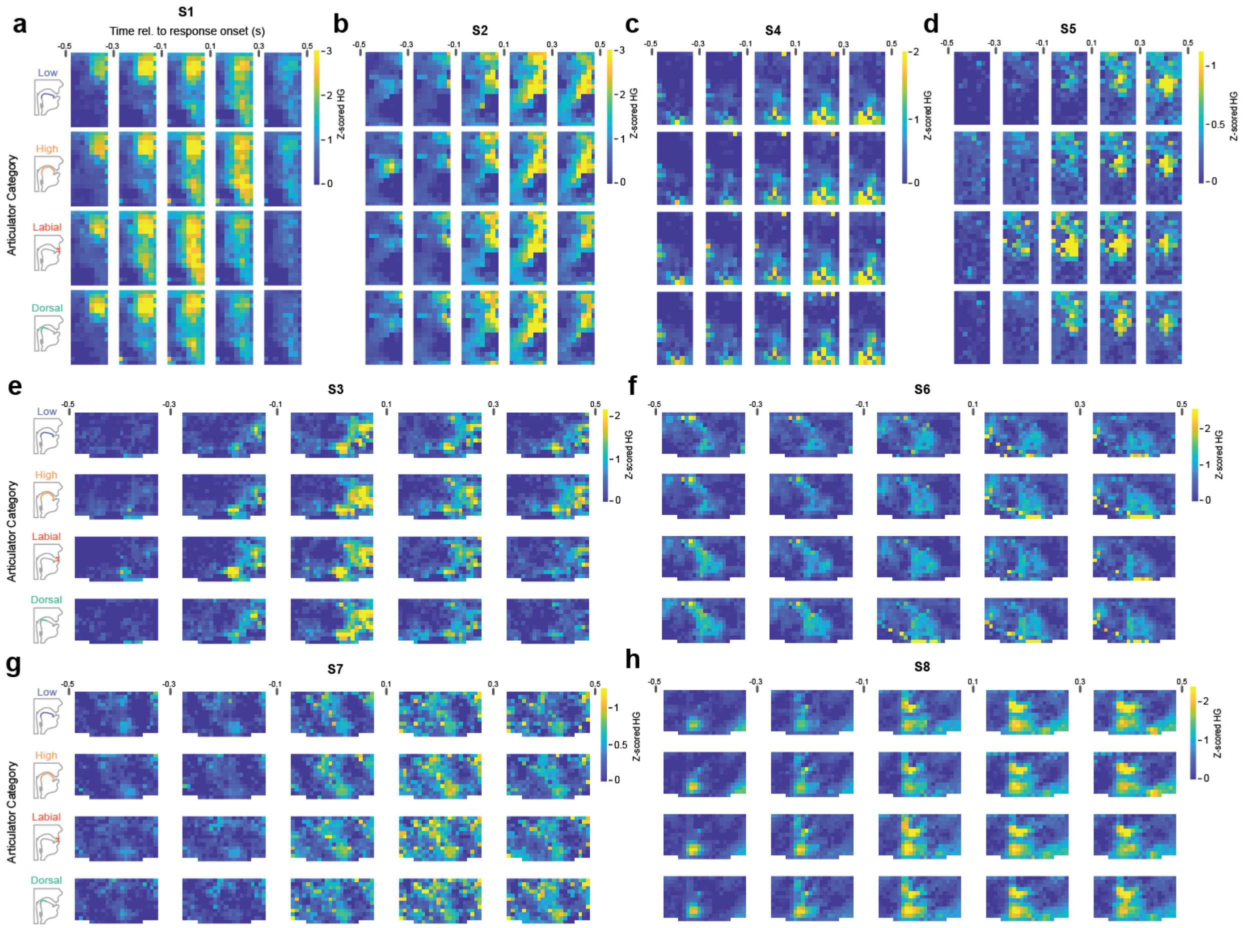


**Figure S4. Spatiotemporal high gamma activations.** Spatiotemporal activation patterns of HG across the µECoG arrays for all patients (S1-S8, **a-h**) grouped by different articulatory gestures. Vocal tract diagrams show visual representation of articulatory category (low, high, labial, dorsal). Note that colorbars reflect different scales per patient to show active regions despite differences in HG z-scores across patients.

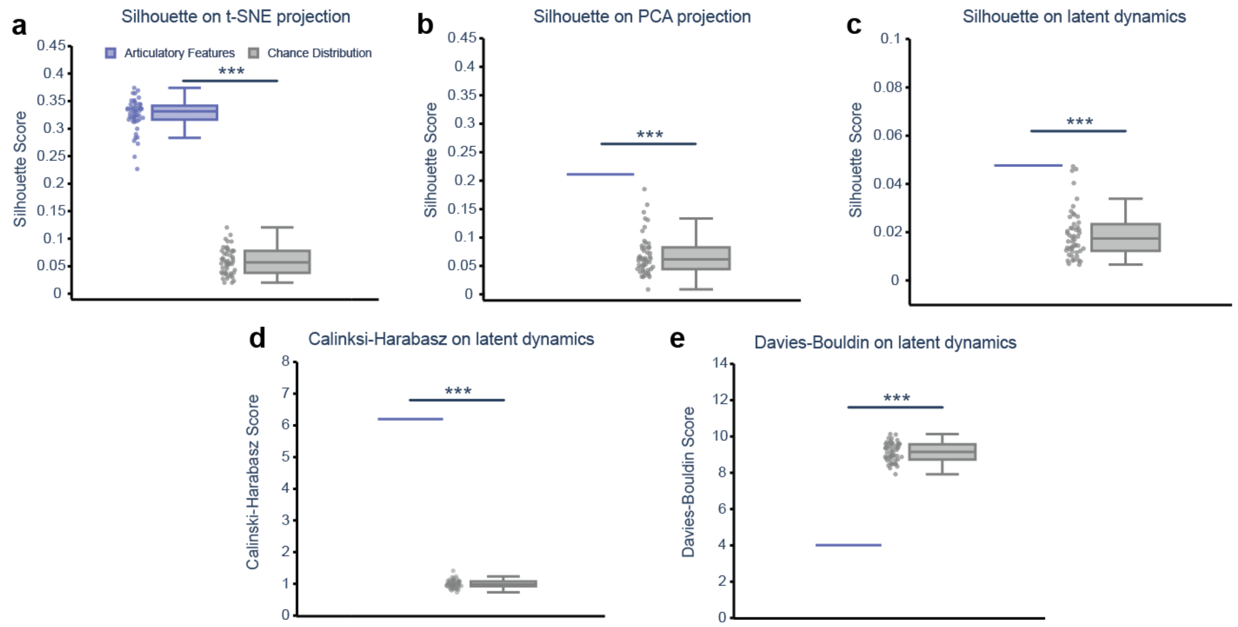


**Figure S5. Clustering of S1 latent dynamics with various metrics. a.** Clustering strength of articulatory features as quantified by the silhouette score on 2D t-SNE-projected latent dynamics (as reported in Fig. 2). Articulatory clustering is stronger than in a shuffled chance distribution (****p* < 0.001, Mann-Whitney U test). As in the main text, the chance distribution is obtained by applying the same clustering quantification after randomly permuting the labels of articulatory features across trials). **b.** Clustering strength as quantified by the silhouette score on 2D PCA-projected latent dynamics. Since PCA results in a deterministic projection, only a single value is generated for the true articulatory features. Articulatory clustering is stronger than in a shuffled chance distribution *(p* < 0.001, one-sample T-test). **c.** Clustering strength directly on the latent dynamics. Articulatory clustering is stronger than in a shuffled chance distribution *(p* < 0.001, one-sample T-test). Due to the relatively high dimensionality of the space (latent components by timepoints), the silhouette score is lower than in **a** and **b**. **d.** Clustering strength as quantified by the Calinski-Harabasz score directly on latent dynamics. Articulatory clustering is stronger than in a shuffled chance distribution *(p* < 0.001, one-sample T-test). **e.** Clustering strength as quantified by the Davies-Bouldin score directly on latent dynamics. A lower score indicates stronger clustering for this metric. Articulatory clustering is stronger than in a shuffled chance distribution *(p* < 0.001, one-sample T-test).

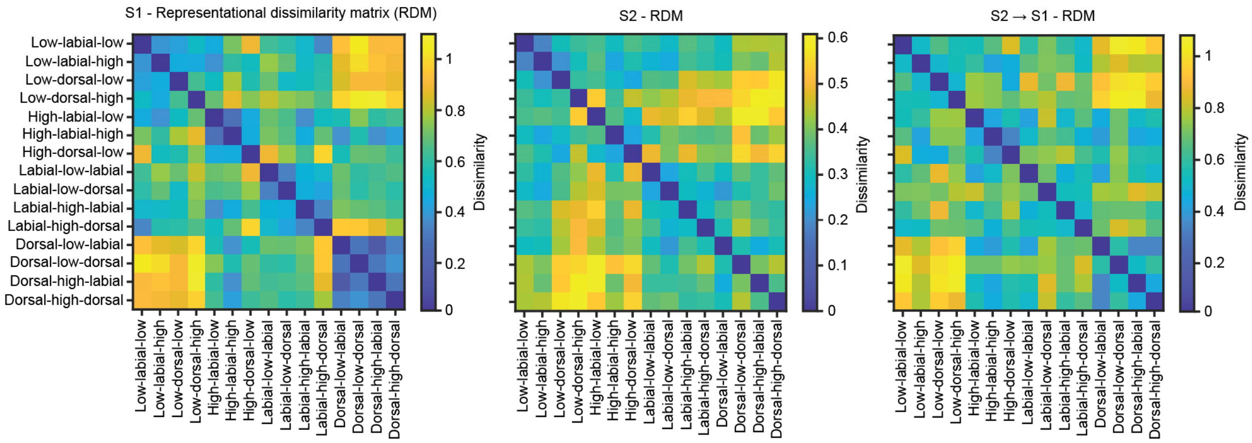


**Figure S6. Example representational dissimilarity matrices.** Representational dissimilarity matrices (RDMs) S1 (left), S2 (middle), and S2 aligned to S1 (right). RDMs are constructed by calculating the dissimilarity (1 – *r*, Pearson correlation) for each pair of experimental conditions. For this analysis, we chose to use labels representing trials as articulatory sequences (three-length sequence of articulators). Notice the difference in experimental structure represented by latent dynamics in S1 and S2. Following alignment of S2 to S1, the RDMs become more visually similar, indicating shared encoding of speech information in aligned latent dynamics.

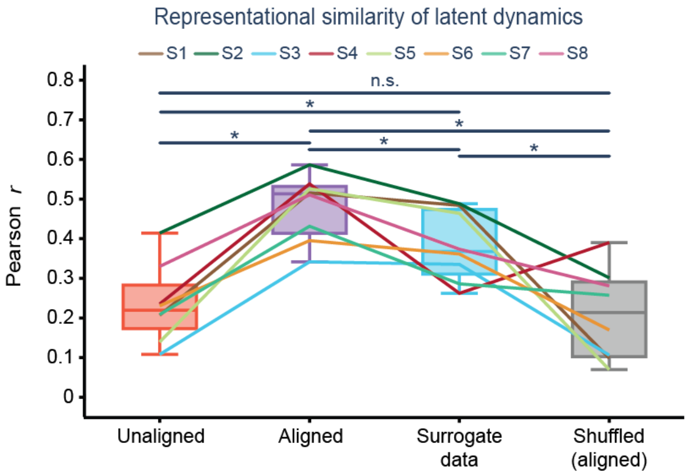


**Figure S7. Speech information improves cross-patient representational similarity.** Cross-patient representational similarity between the latent dynamics of each target patient and all source patients. Representational similarity is calculated as the Pearson correlation between RDMs across patients. The first two columns are as reported in Fig. 2f. In the third column, the representational similarity analysis is repeated using surrogate data (see Methods) that preserves neural structure but is random with respect to speech. In the fourth column, representational similarity on true aligned data using shuffled labels, as a negative control. We find all significant differences (**p* < 0.05, FDR-corrected Wilcoxon signed-rank test) except for between unaligned and shuffled conditions (*p*=0.31, FDR-corrected Wilcoxon signed-rank test).

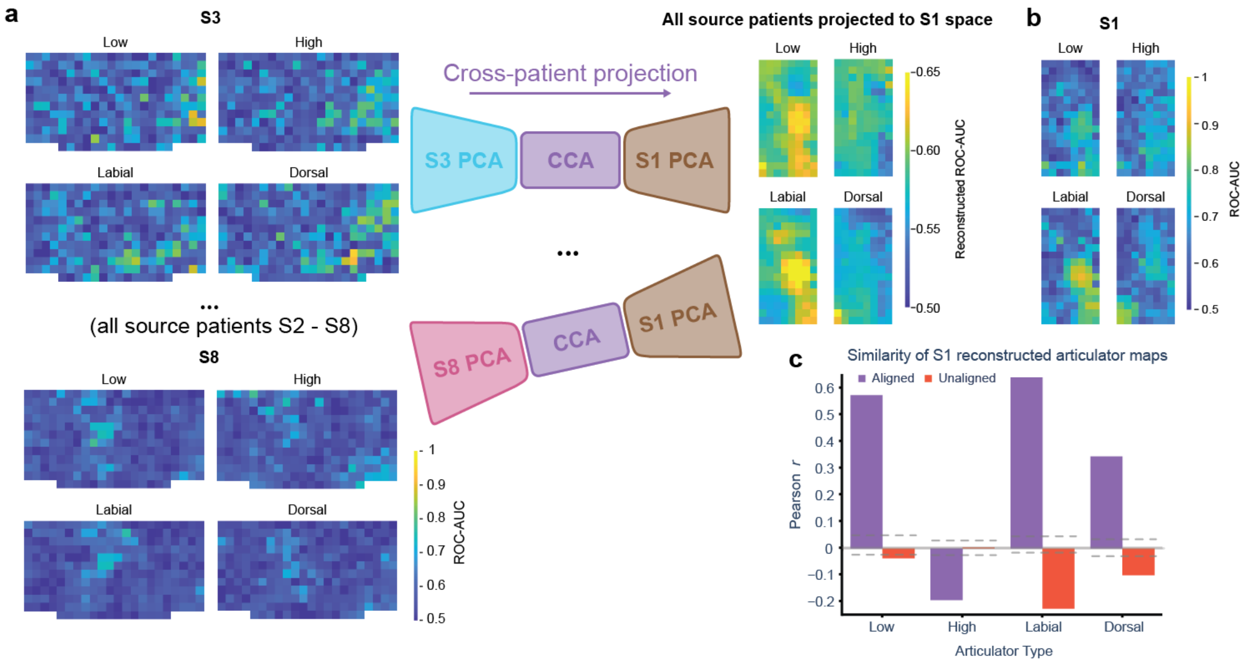


**Figure S8. Cross-patient projection averaged across source patients. a.** Cross-patient projection, as in Fig. 3, but across multiple source patients. The articulator maps of all patients S2-S8 (left, S3 and S8 shown here) are projected to S1's electrode-space (see Methods) and the resultant reconstructed maps are averaged across source patients within articulator type (right). This method considers reconstructed speech information across all source patients instead of a singular source patient as shown in the Fig. 3b. **b.** Ground-truth S1 articulator maps for reference (as reported in Fig. 3a). **c.** Similarity between source-patient-averaged reconstructed articulator maps and ground truth S1 articulator maps, as measured by Pearson correlation. Horizontal dashed lines indicate an interval of trivial correlation values, represented by 95% confidence intervals on correlation values from a null distribution of reconstructions performed with permuted CCA weights ($n$ = 100). All articulator types except "high" show improved similarity when compared to reconstructions of S1 articulator maps by only S3 (as in Fig. 3c, left), while the "high" type shows misalignment to the target articulator map, as shown by its non-trivial anti-correlation.

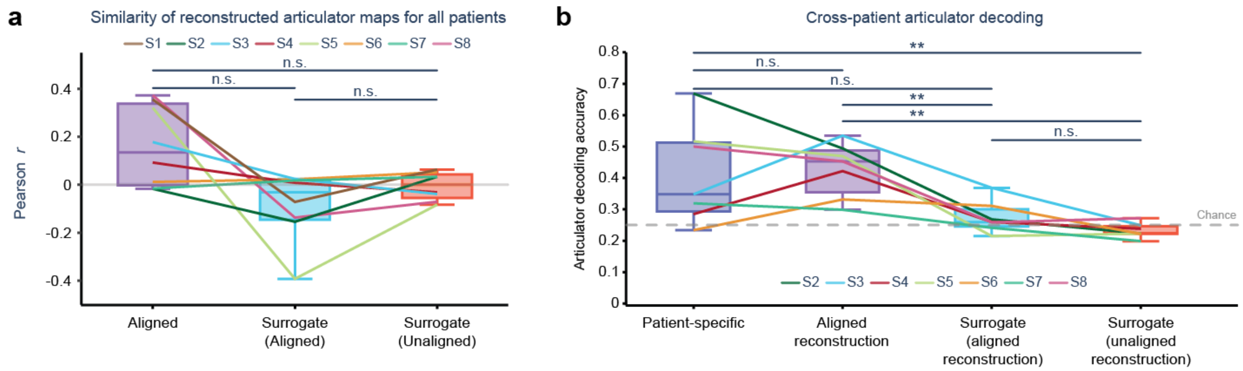


**Figure S9*. Cross-patient projection on surrogate data.*** Repetition of cross-patient projection experiment (Fig. 3) while incorporating surrogate data control (see Methods). **a.** While there is no significant difference between reconstructed articulator map similarity with true or surrogate data ($p = 0.11$, FDR-corrected Wilcoxon signed-rank test), we do notice a relatively large drop in mean from true aligned to surrogate data. **b.** Cross-patient articulator decoding with surrogate data. We see a significant reduction in decoding accuracy between aligned reconstructions with true and surrogate data (**$p < 0.01$, FDR-corrected Wilcoxon signed-rank test).

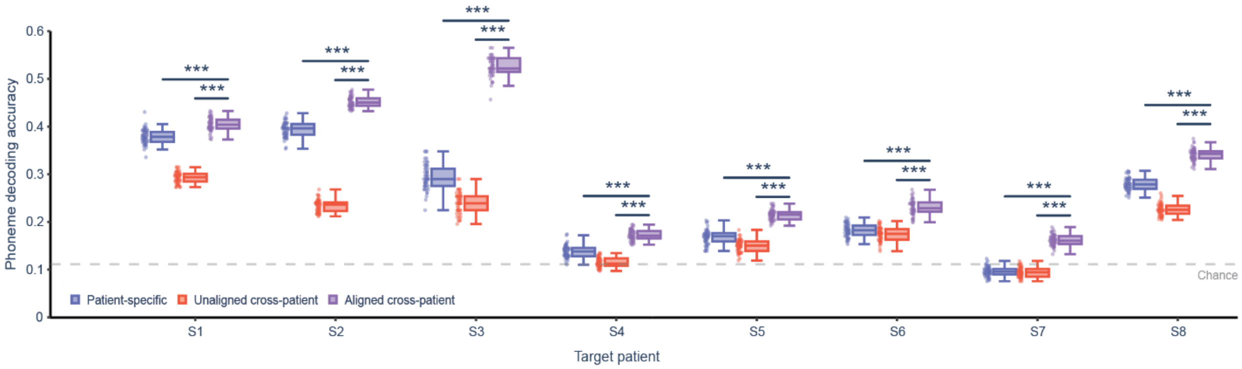


**Figure S10. Cross-patient phoneme decoding by target patient.** Cross-patient phoneme decoding for each target patient separately. Aligned, cross-patient data results in significantly higher phoneme decoding than both unaligned and patient-specific data for all patients (****p < 0.001*, post-hoc Tukey HSD test following one-way ANOVA within target patient). The gray dotted line indicates chance levels of decoding.

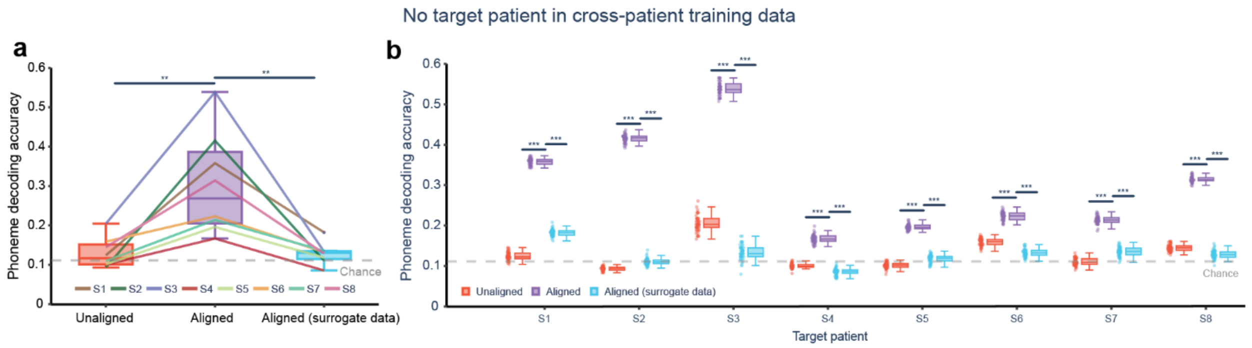


**Figure S11*. Cross-patient phoneme decoding on non-target patient data.*** As in Fig. 4 and Fig. S7, but data from the target patient is excluded from the training set of the decoding model (i.e. target patient data is only used to align other patients to the target space). The gray dotted line indicates chance levels of decoding. **a,** Without alignment, data from other patients results in chance levels of phoneme decoding. With alignment, we see significantly higher phoneme decoding accuracies across all patients (***p < 0.01*, repeated measures ANOVA and follow-up FDR-corrected paired t-tests). To confirm that the increased decoding accuracies from alignment were not artificially induced by the alignment procedure, surrogate data was generated that preserved second-order statistics of neural data from each patient but was otherwise random (see Methods) and was aligned to a non-surrogate target patient prior to phoneme decoding. This aligned surrogate data achieved approximately chance levels of decoding and had significantly lower phoneme decoding accuracy than the aligned non-surrogate context, validating the aligned decoding approach. **b,** As in **a** but for each target patient individually. Alignment of true neural data results in higher accuracies than unaligned and aligned surrogate data for all target patients, even without the inclusion of the target patient in the training data (****p<0.001,* post-hoc Tukey test following one-way ANOVA within target patient).

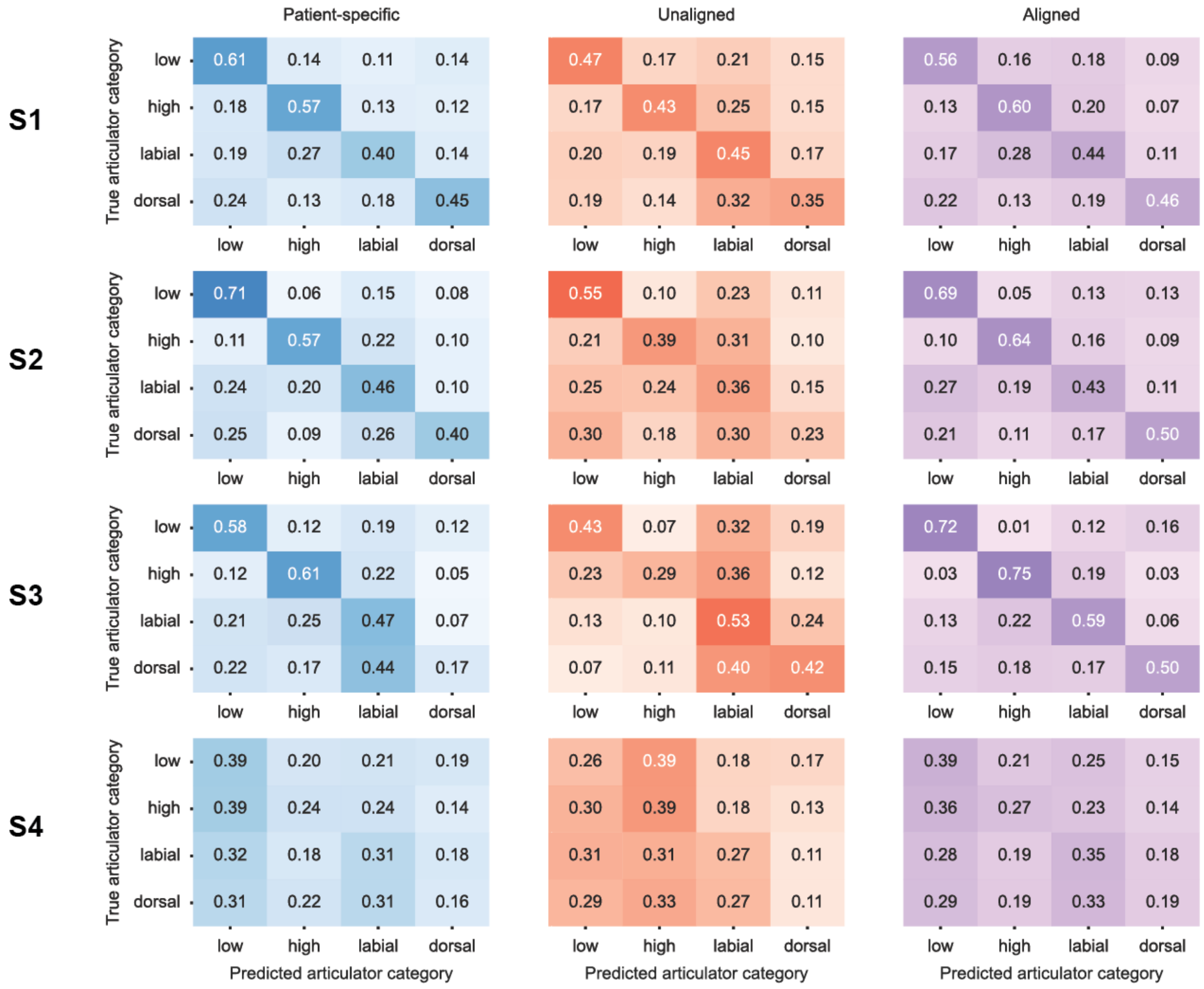


**Figure S12. Articulatory decoding confusion matrices (S1-S4).** Confusion matrices are normalized such that each row sums to 1. Confusion matrices are aggregated over 50 iterations of decoding models within each patient and decoding context.

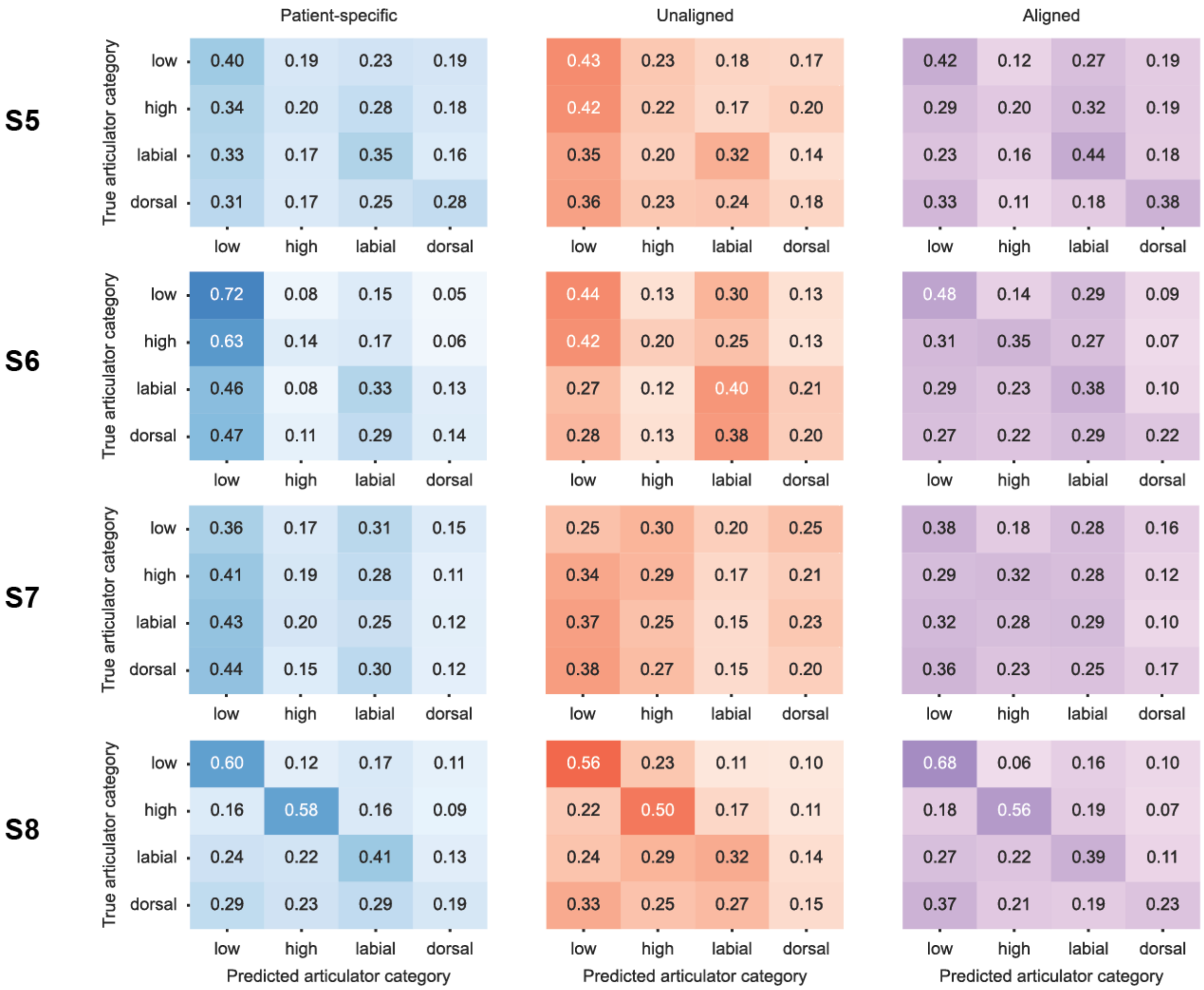


**Figure S13. Articulatory decoding confusion matrices (S5-S8).** Confusion matrices are normalized such that each row sums to 1. Confusion matrices are aggregated over 50 iterations of decoding models within each patient and decoding context.

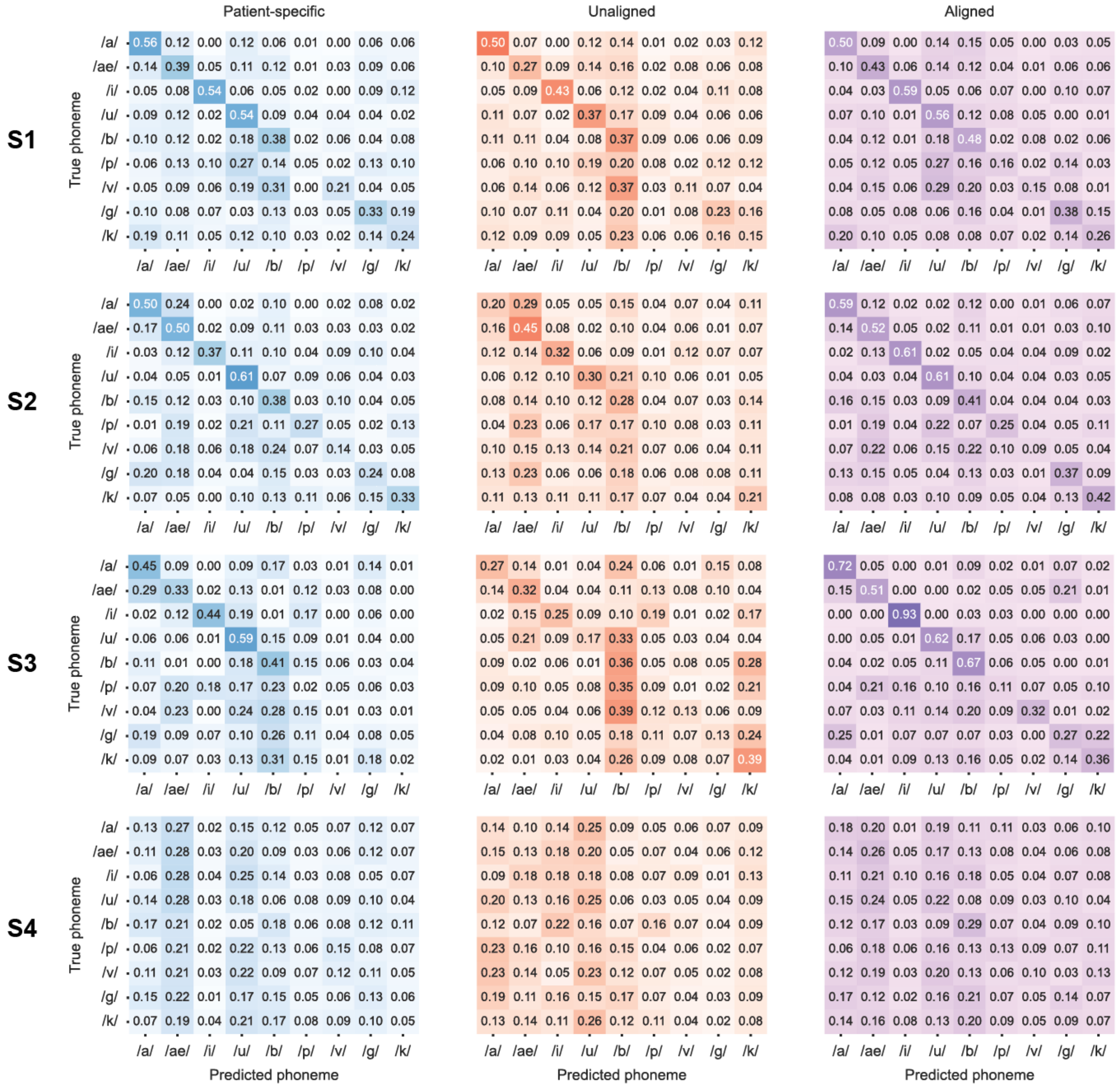


**Figure S14. Phoneme decoding confusion matrices (S1-S4).** Confusion matrices are normalized such that each row sums to 1. Confusion matrices are aggregated over 50 iterations of decoding models within each patient and decoding context.

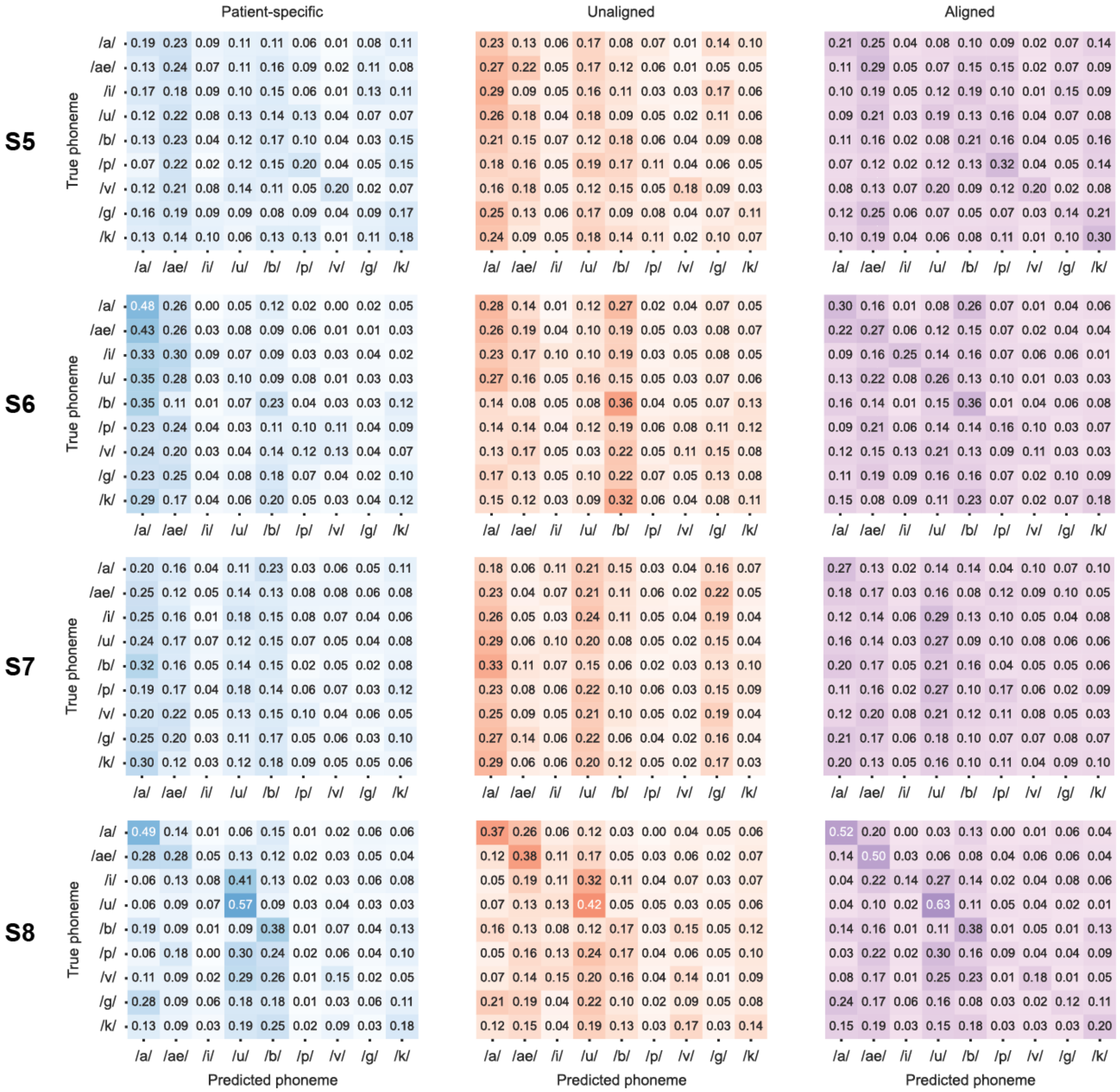


**Figure S15. Phoneme decoding confusion matrices (S5-S8).** Confusion matrices are normalized such that each row sums to 1. Confusion matrices are aggregated over 50 iterations of decoding models within each patient and decoding context.

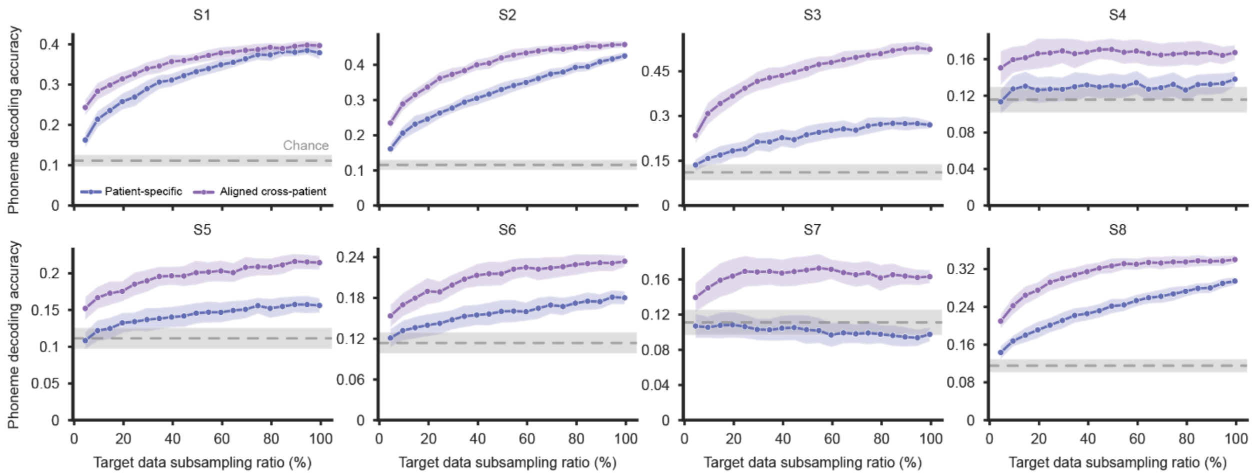


**Figure S16*. Target patient subsampling by patient.*** As reported in Fig. 4c, but plotted separately for each target patient. Patient-specific and aligned cross-patient decoding is performed as the amount of data from the target patient (used for both alignment and decoding) is subsampled. Values are reported every 5% of target data. The gray dotted line indicates the mean accuracy of chance decoding, with error shown as ±1 standard deviation.

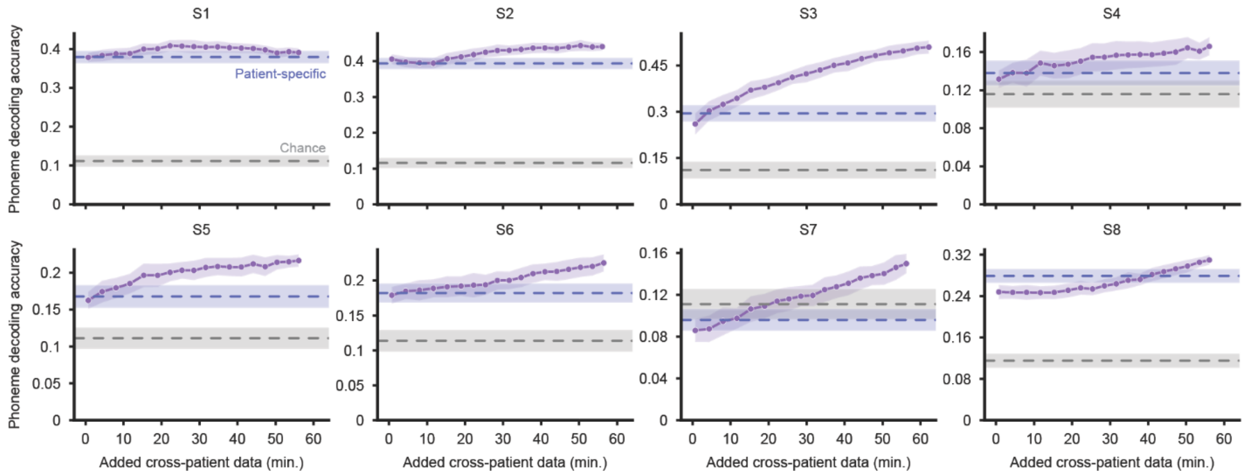


**Figure S17*.* Cross-patient trial subsampling by patient.** As reported in Fig. 4d (only S5), but plotted separately for each target patient. Aligned cross-patient decoding is performed as the amount of cross-patient data is subsampled. The gray dotted line indicates the mean accuracy of chance decoding, with error shown as ±1 standard deviation. The blue dotted line indicates the mean patient-specific decoding accuracy for each target patient, with error shown as ±1 standard deviation.

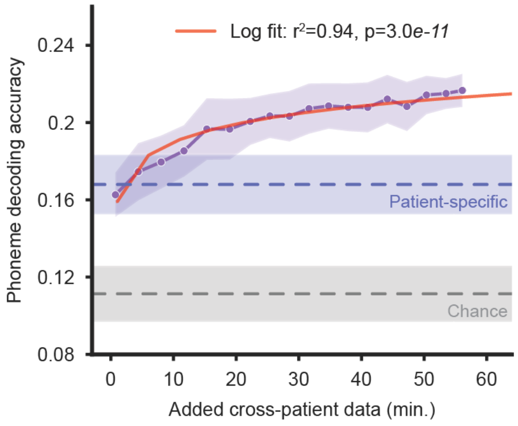


**Figure S18. Log fit to S5 cross-patient subsampling data.** As reported in Fig. 4d, but with a log fit to S5's phoneme decoding accuracy vs. cross-patient trials included data (performed as a linear fit to log-transformed data, $p$ = 3e-11). The log fit was used to confirm diminishing gains in decoding accuracy for the SVM model as more cross-patient data is added. The log-fit was well-matched to the data with $r^2$ = 0.94. The gray dotted line indicates the mean accuracy of chance decoding, with error shown as ±1 standard deviation. The blue dotted line indicates the mean patient-specific decoding accuracy for each target patient, with error shown as ±1 standard deviation.

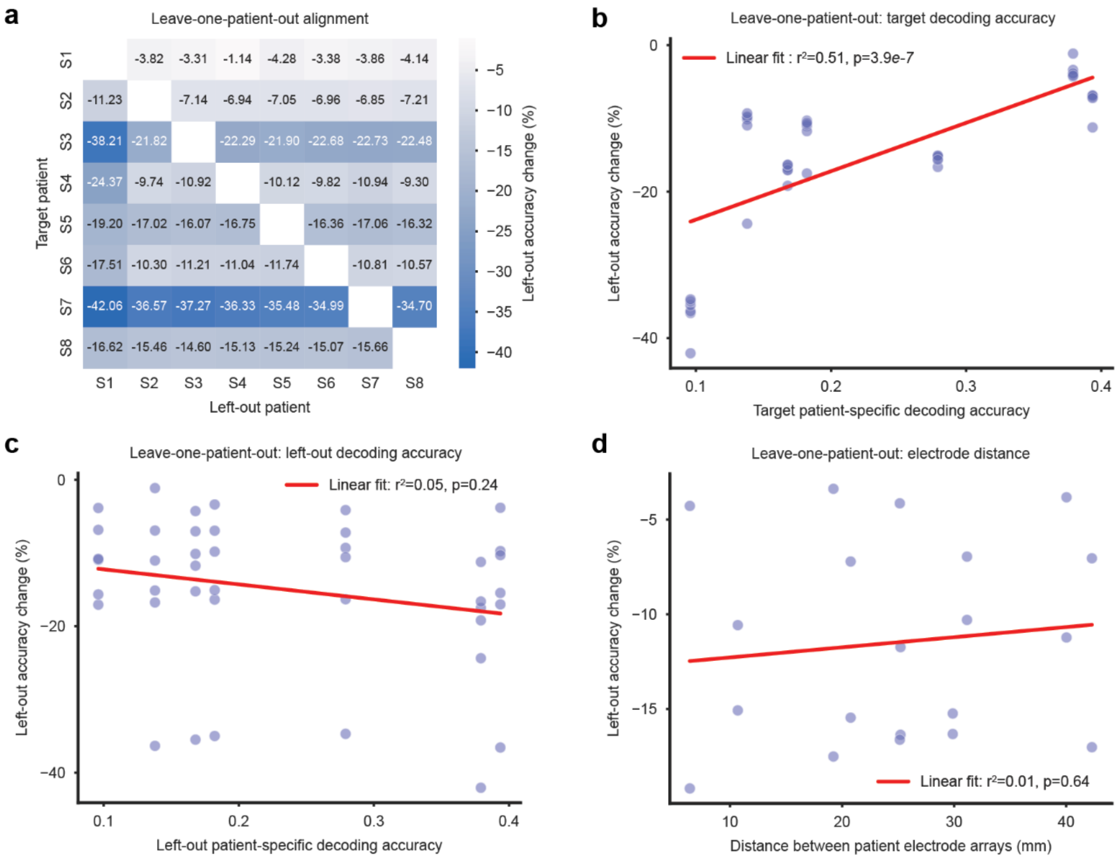


**Figure S19. Leave-one-out inter-patient variability.** Pairwise patient effects quantified by parametric leaving-out of patient from cross-patient training set. Left-out accuracy change is quantified by the percent change between target patient phoneme decoding with all source patients and with all source patients but the parametrically removed one. **a.** Heatmap of left-out accuracy results by target patient and left-out patient. **b.** We found a significant, positive relationship between target patient-specific decoding accuracy and left-out accuracy change, indicating that, target patients that perform better have less reliance on any single cross-patient dataset. We found no significant effect of the patient-specific accuracy of the left-out patient (**c**) or distance between electrodes in neuroanatomical space (**d**).

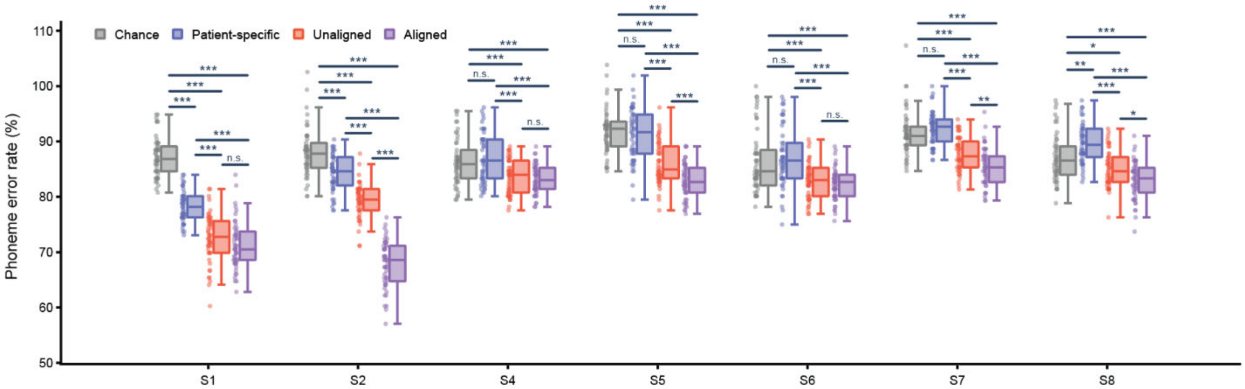


**Figure S20*. CTC-RNN decoding by target patient.*** Note: This plot shows error rate, so lower phoneme error rate (PER) is better performance. CTC-RNN decoding for each patient and decoding context separately. Patient-specific PERs are either at or only slightly below chance for most patients. Unaligned PERs are significantly lower than patient-specific in all cases, surprisingly. We see significantly lower error rates in aligned cases than unaligned cases for most patients, though not all. Aligned PERs are significantly lower than patient-specific PERs. Within patients, we performed a one-way ANOVA and follow-up Tukey HSD tests. Displayed p-values are $^{*}p < 0.05$, $^{**}p < 0.01$, $^{***}p < 0.001$)

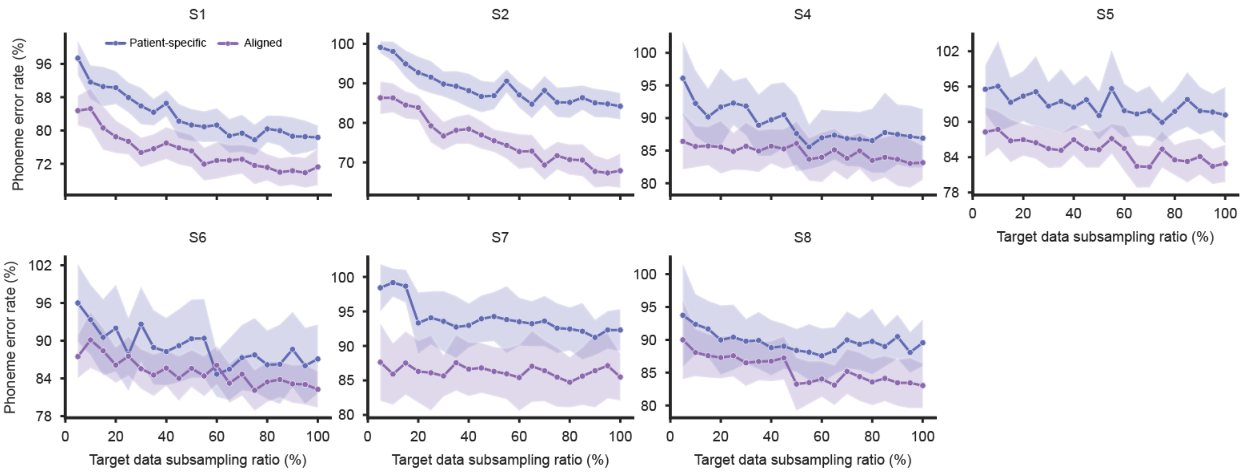


**Figure S21. CTC-RNN target patient subsampling by patient.** As reported in Fig. 5c, but plotted separately for each target patient. Patient-specific and aligned cross-patient decoding is performed as the amount of data from the target patient (used for both alignment and decoding) is subsampled. Values are reported every 5% of target data.

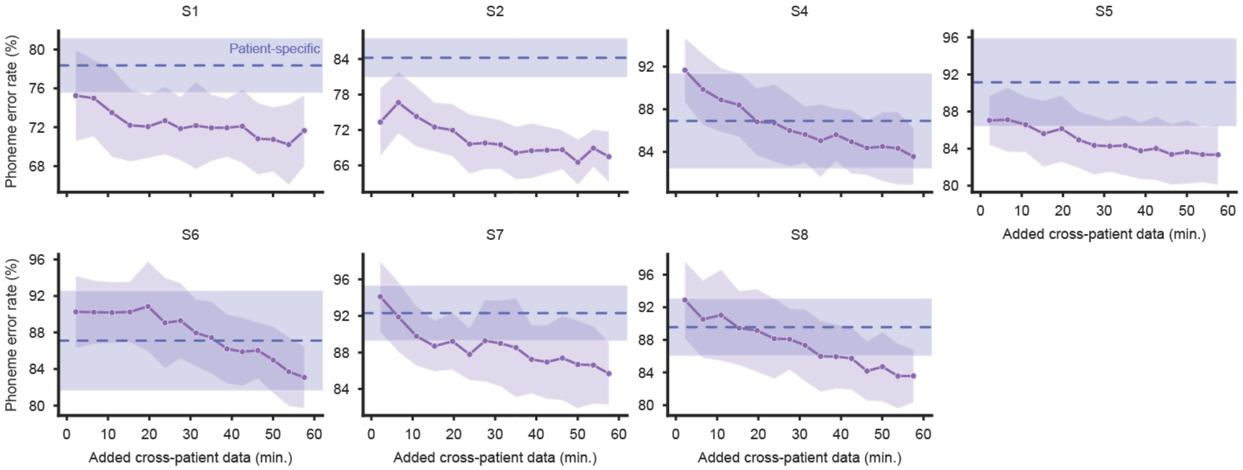


**Figure S22. CTC-RNN target patient subsampling by patient.** As reported in Fig. 5d, but plotted separately for each target patient. Aligned cross-patient decoding is performed as the amount of cross-patient data is subsampled. The blue dotted line indicates the mean patient-specific decoding accuracy for each target patient, with error shown as ±1 standard deviation.

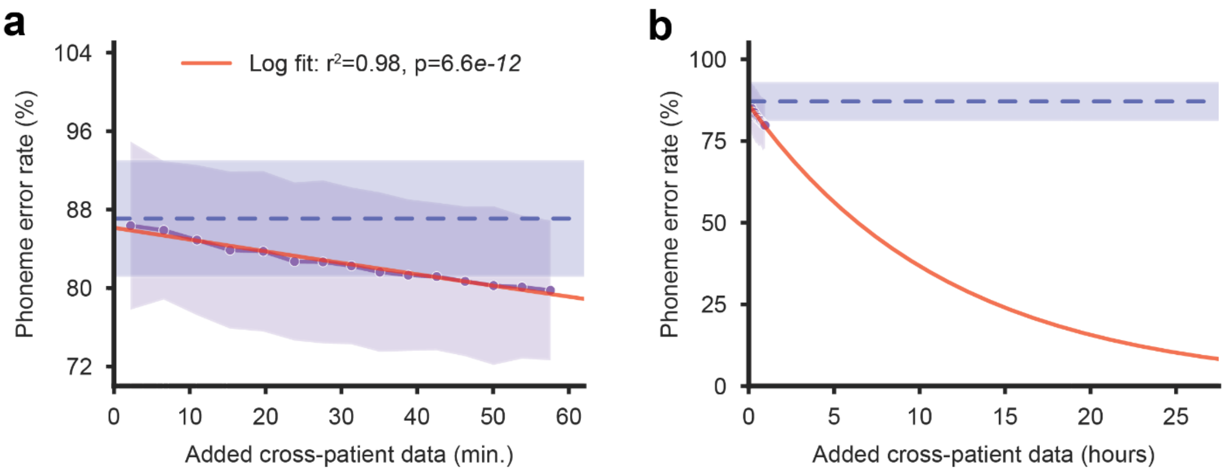


**Figure S23*. Log fit to CTC-RNN cross-patient subsampling data.* a.** As reported in Fig. 5d, but with a log fit to PER vs. amount of cross-patient data included data (performed as a linear fit to log-transformed data, $p$ = 6.6e-12). The log-fit was well-matched to the data with $r^2$ = 0.98. **b.** The log fit was used to project cross-patient data amounts to data amounts presented in state-of-the-art ECoG speech BCIs. The blue dotted line indicates the mean patient-specific decoding accuracy for each target patient, with error shown as ±1 standard deviation.

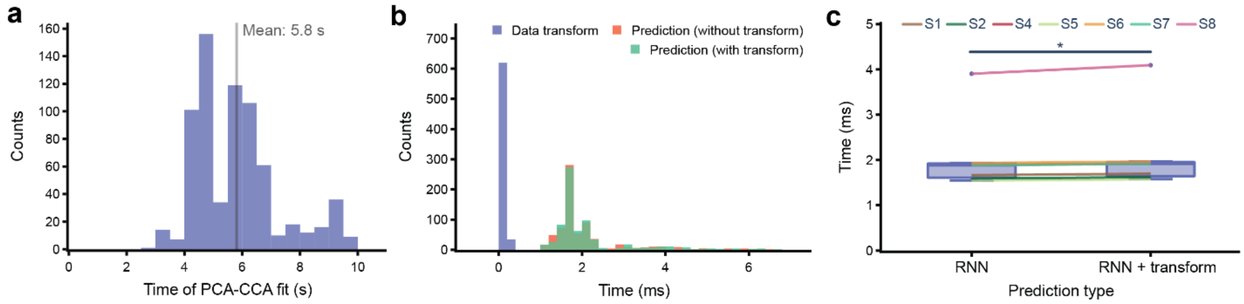


**Figure S24*. *Quantification of aligned CTC-RNN latencies. a.** Latency of PCA-CCA fit to learn decomposition and alignment on the entire training set. This is only a one-time calculation following the collection of a small amount of training/calibration data. We found a mean fit time of ~6 seconds and a max of ~ 10 seconds, both of which negligible one-time delays compared to the ~6 minutes of patient-specific data collection. **b.** Latency of CTC-RNN model predictions, both with a linear input layer attached and without. The additional linear input layer is necessary if aligning real-time collected data from a target patient instead of aligning previously collected data to the space of the target patient. **c.** Comparison of RNN prediction latencies with and without the added linear transform for all patients. While we do find a significant difference between prediction types (*$p < 0.05$, Wilcoxon signed-rank test), it is only a small difference of ~ 0.05 ms. As such, both mean prediction times of 2.06 ms (RNN) and 2.11 ms (RNN + transform) are well under our latency budget of 80 ms, indicating feasible implementation.

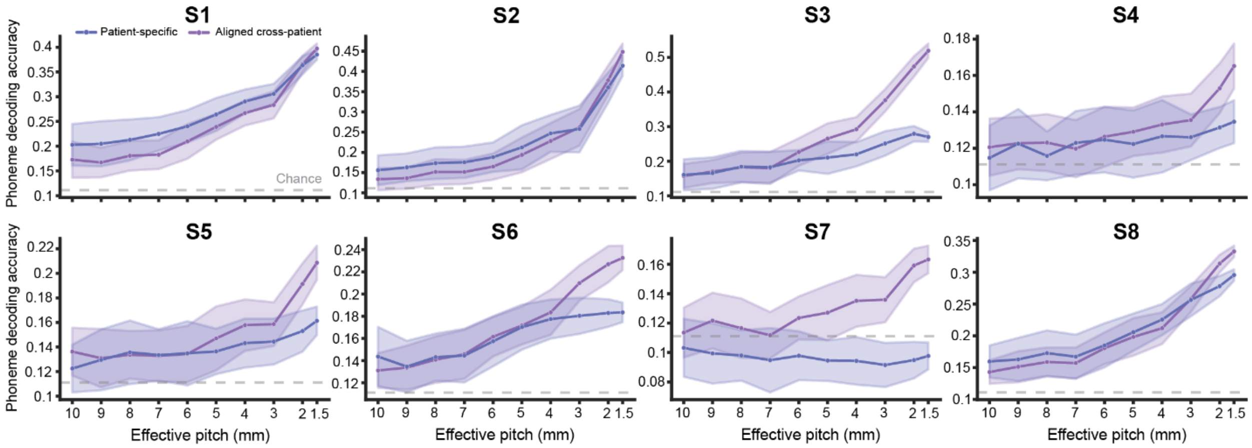


**Figure S25. Pitch-subsampled phoneme decoding by patient.** As reported in Fig. 6a, but plotted separately for each target patient. Patient-specific and aligned cross-patient decoding is performed with Poisson-disk-subsampled electrode data to simulate different electrode array densities. The gray dotted line indicates chance levels of decoding.

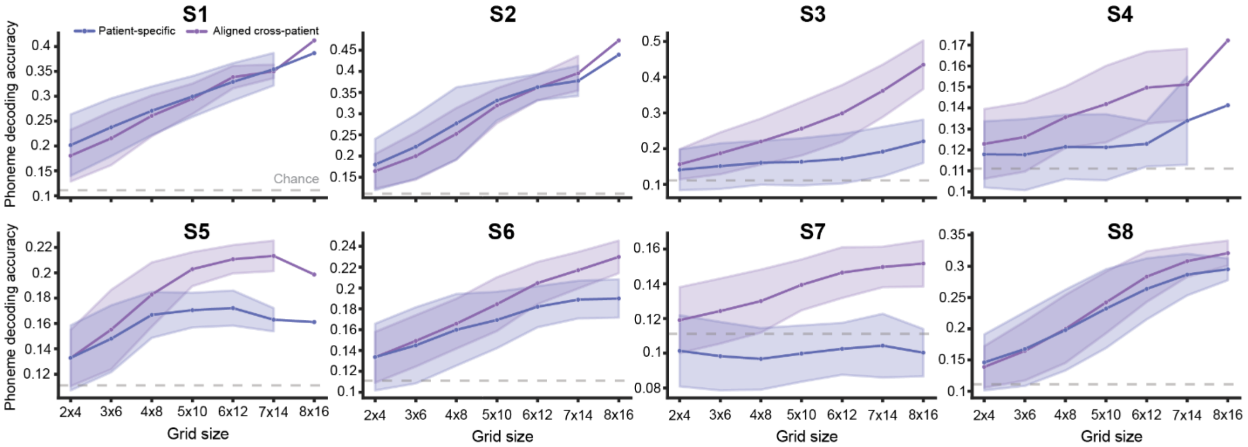


**Figure S26. Grid-subsampled phoneme decoding by patient.** As reported in Fig. 6b, but plotted separately for each target patient. Patient-specific and aligned cross-patient decoding is performed with data from subsampled grids of electrodes to simulate different electrode array coverages. The gray dotted line indicates chance levels of decoding. For patients with 128-channel arrays (S1, S2, S4, S5), there is a singular point for the 8x16 grid size since that encompasses the entire electrode array.

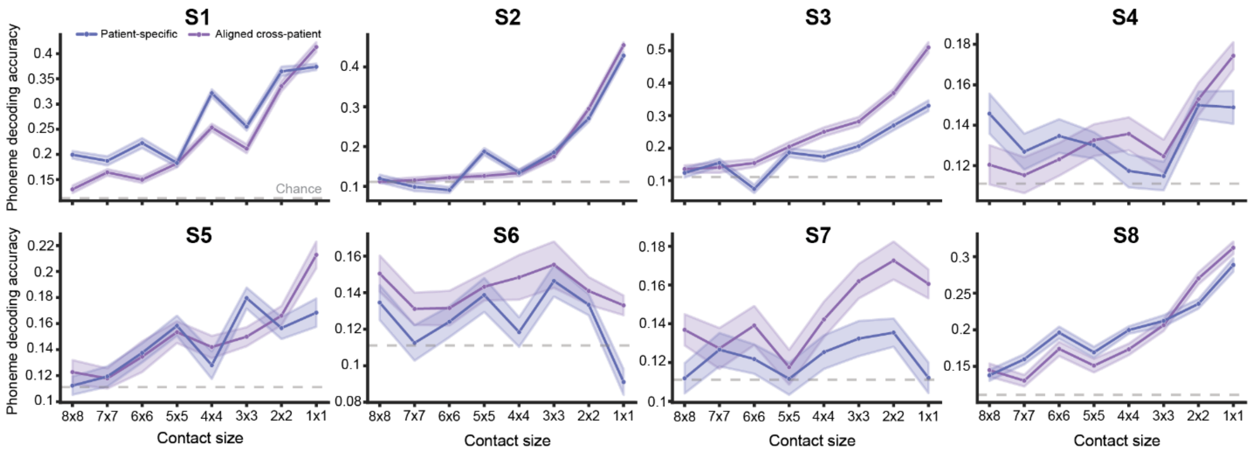


**Figure S27.** **Spatially-averaged phoneme decoding by patient.** As reported in Fig. 6c, but plotted separately for each target patient. Patient-specific and aligned cross-patient decoding is performed with HG data extracted from averaged electrode contacts of various sizes to simulate recording with larger electrode contacts. The gray dotted line indicates chance levels of decoding.
